# Rapid and efficient adaptation of the dTAG system in mammalian development reveals stage specific requirements of NELF

**DOI:** 10.1101/2021.11.30.470581

**Authors:** Abderhman Abuhashem, Anna-Katerina Hadjantonakis

## Abstract

Targeted protein degradation methods offer a unique avenue to assess a protein’s function in a variety of model systems. Recently, these approaches have been applied to mammalian cell culture models, enabling unprecedented temporal control of protein function. However, the efficacy of these systems at the tissue and organismal levels in vivo is not well established. Here, we tested the functionality of the degradation tag (dTAG) degron system in mammalian development. We generated a homozygous knock-in mouse with a FKBP^F36V^ tag fused to Negative elongation factor b (Nelfb) locus, a ubiquitously expressed protein regulator of transcription. In the first validation of targeted endogenous protein degradation across mammalian development, we demonstrate that irrespective of the route of administration the dTAG system is safe, rapid, and efficient in embryos from the zygote to midgestation stages. Additionally, acute early depletion of NELFB revealed a specific role in zygote-to-2-cell development and Zygotic Genome Activation (ZGA).

**HIGHLIGHTS:**

- genetically engineered mouse model harboring a FKBP^F36V^ knock-in to evaluate kinetics and efficacy of the dTAG degron system in vivo
- system is non-toxic, and allows acute and efficient degradation of a FKBP^F36V^- tagged endogenous protein during in utero embryo development
- system nables fine temporal degradation and reversibility of depletion across embryonic stages
- stage-specific depletion reveals a role for NELFB during mouse ZGA

## INTRODUCTION

Molecular perturbation methods have been evolving rapidly over the last few decades and provide an invaluable tool to interrogate gene function across biological contexts. Broadly, these methods can be categorized based on their point of action: DNA, RNA, or protein. The recent revolution in CRISPR systems enormously expanded the DNA/RNA targeting toolbox (Pickar-Oliver and Gersbach, 2019). However, because proteins are the functional biological units in the majority of contexts, targeting DNA/RNA poses several limitations. These include slow onset of effect and the possibility of adaptation, irreversibility when DNA is modified, and biological plasticity from alternative splicing (Smits et al., 2019; Verma et al., 2020; Wu et al., 2020). These limitations hinder experimental design and ultimately appropriate interpretation of protein functions in scenarios that require acute perturbation and/or reversibility.

Direct protein perturbation is an attractive alternative that may offer superior temporal resolution to DNA/RNA targeting methods (Wu et al., 2020). While small molecules can be used to target some proteins, the druggable proteome currently represents a small subfraction of proteins in a cell (Chamberlain et al., 2019). Recent development of degron approaches, such as the auxin-inducible degron (AID) and degradation tag (dTAG) systems offer the possibility of targeting any protein in a cell in a rapid, inducible, and reversable manner (Verma et al., 2020). These systems rely on genetic fusion of the gene of interest to a small tag. Upon introduction of a small molecule that binds specifically both the tag and ubiquitination proteins, the tagged protein can be directed for proteasomal degradation. These system have enabled discoveries in cell models that were not previously possible (Jaeger and Winter, 2021). Notably, the dTAG system exclusively relies on the mammalian endogenous proteasomal degradation machinery, while the AID system requires the expression of a exogenous plant-based factor to activate the system (Nabet et al., 2018; Nishimura et al., 2009). Despite their promise, it is unclear how safe or functional these systems are in vivo, particularly during mammalian development.

Early mammalian development poses a unique challenge for interrogation of protein function. During pre-implantation embryo development, maternally deposited transcripts can mask early loss of function (knockout) phenotypes, often necessitating generation of maternal/zygotic mutants through complex and imperfect Cre/loxP genetic systems that knockout both maternal and zygotic genes (Sun et al., 2008). Early post-implantation, Cre-estrogen receptor (Cre-ER) systems can offer some spatio-temporal specificity, but can result in mosaic recombination across and within embryos, while tamoxifen injections can have teratogenic effects on embryos, as well as leading to implantation failure during these stages (Pimeisl et al., 2013; Savery et al., 2020; Ved et al., 2019). Thus, there is a need for a safe, efficient, rapid, inducible, and reversable perturbation system to probe gene functions in complex mammalian systems. Ideally, such a system would also enable iterative and/or successive interrogation of gene function at a variety of developmental stages.

Here, leveraging recent observations in mammalian cells and compelling safety and specificity profile, we sought to implement the dTAG system in mammalian embryos. We tagged Negative Elongation Factor-b (NELFB) with FKBP^F36V^ in mouse embryonic stem cells (mESCs). Nelfb is widely expressed across mouse development, enabling validation of the dTAG system at a wide range of stages (Nowotschin et al., 2019; Pijuan-Sala et al., 2019). Then, we used these mESCs to generate germline-transmitting mice. Resulting F1 animals heterozygous for the *Nelfb-FKBP^F36V^* allele were subsequently used to generate a homozygous mouse with this allele, demonstrating the fusion to be safe. Using this *Nelfb-FKBP^F36V^* allele, we demonstrated that the dTAG system provides highly efficient acute protein degradation across pre- and post- implantation embryonic stages (E0.5 – E9.5). We found that dTAG small molecules dTAG-13 and dTAGv-1 are developmentally neutral and safe for use in in vitro embryo culture and via intraperitoneal injection into pregnant adult females. The system achieves maximum efficiency within one hour in ex utero cultured pre-implantation embryos, and within ∼4 hours in vivo in post-implantation embryos. We utilized the system to uncover a novel role for NELFB in pre-implantation development, specifically in zygotic genome activation (ZGA). We found that NELFB attenuates the initiation of major ZGA. To our knowledge, this is the first successful adaptation of a degron system to perturb an endogenous protein in a mouse model and during embryonic stages.

## DESIGN

We considered several degron systems for application in the context of mammalian embryonic development including the AID, dTAG, and TRIM-away systems (Clift et al., 2017; Nabet et al., 2018; Nishimura et al., 2009). While the TRIM-away system does not require gene editing as it relies on antibodies to identify target proteins, it can be technically challenging depending on the sub-cellular localization of the target protein and consequently antibodies, which are not cell permeable (Clift et al., 2017). Both the AID and dTAG systems offer comparable advantages in terms of acute degradation of a genetically edited tagged target protein (Nabet et al., 2018; Nishimura et al., 2009). The second iteration of the AID system, AID2, overcame significant challenges of basal degradation and has been shown to be effective in degrading a tagged transgene reporter in vivo in mice (Nishimura et al., 2020; Yesbolatova et al., 2020). However, auxin derivatives have been shown to be bioactive and can potentially affect embryonic development (Nishimura et al., 2020). Furthermore, all auxin systems require constitutive exogenous expression of a plant-based accessory protein (e.g. OsTIR1). By contrast, the dTAG system only requires editing a locus of interest to introduce a FKBP^F36V^ tag. Additionally, several recently developed dTAG molecules (e.g. dTAG-13 and dTAGv-1) can induce degradation at nanomolar concentrations and do not appear to induce biological activity in the absence of the FKBP^F36V^ tag (Nabet et al., 2018, 2020). Depending on the dTAG molecule used, either CRBN or a VHL-mediated proteasomal degradation is engaged, which can theoretically expand the functionality of the system to more cell types.

NELFB regulates a critical step in transcription, promoter-proximal RNA Pol II pausing. As a result, it is present across mammalian development and tissues (Nowotschin et al., 2019; Pijuan-Sala et al., 2019). It therefore represents an ideal choice of locus to assess the dTAG system across a variety of stages and tissues. Furthermore, the role of NELFB has not been assessed at early developmental stages and a recent study suggested a dynamic role for NELFB at the zygote/2-cell stages (Liu et al., 2020). These considerations motivated us to design a successful mouse model with degradable NELFB and to test its function at pre-implantation stages.

## RESULTS

### Generation of *Nelfb^dTAG^ mouse to* test the dTAG system in vivo

To assess use of the dTAG system in vivo, we sought to generate a mouse model harboring an endogenously tagged protein. To do so, we utilized CRISPR/Cas9 and homology-directed repair (HDR) to insert FKBP^F36V^ in-frame at the C-terminus of NELFB by targeting the final coding exon 5’ of the STOP and 3’ UTR in mESCs (Figure S1A)(Nabet et al., 2018; Ran et al., 2013). Structural studies have shown that the c-terminus of NELFB is not involved in critical interactions of the protein (Vos et al., 2018a). The edited allele also included two HA tags to facilitate detection. mESC clones with correct integration of the tag at the Nelfb locus were identified by PCR (Figure S1B). To ensure that the tagged allele, *Nelfb^dTAG^, was prop*erly expressed and translated, we performed immunofluorescence against the HA tag in a heterozygous clone, which showed robust nuclear staining as expected for NELFB (Figure 1B). Western blot analysis using an antibody against NELFB showed two bands separated by ∼13kD, corresponding to the molecular weight of FKBP^F36V^ (Figure 1C). The bands show similar intensities, suggesting that NELFB-FKBP^F36V^ is stable in the absence of degradation promoting small molecules.

**Figure 1.**
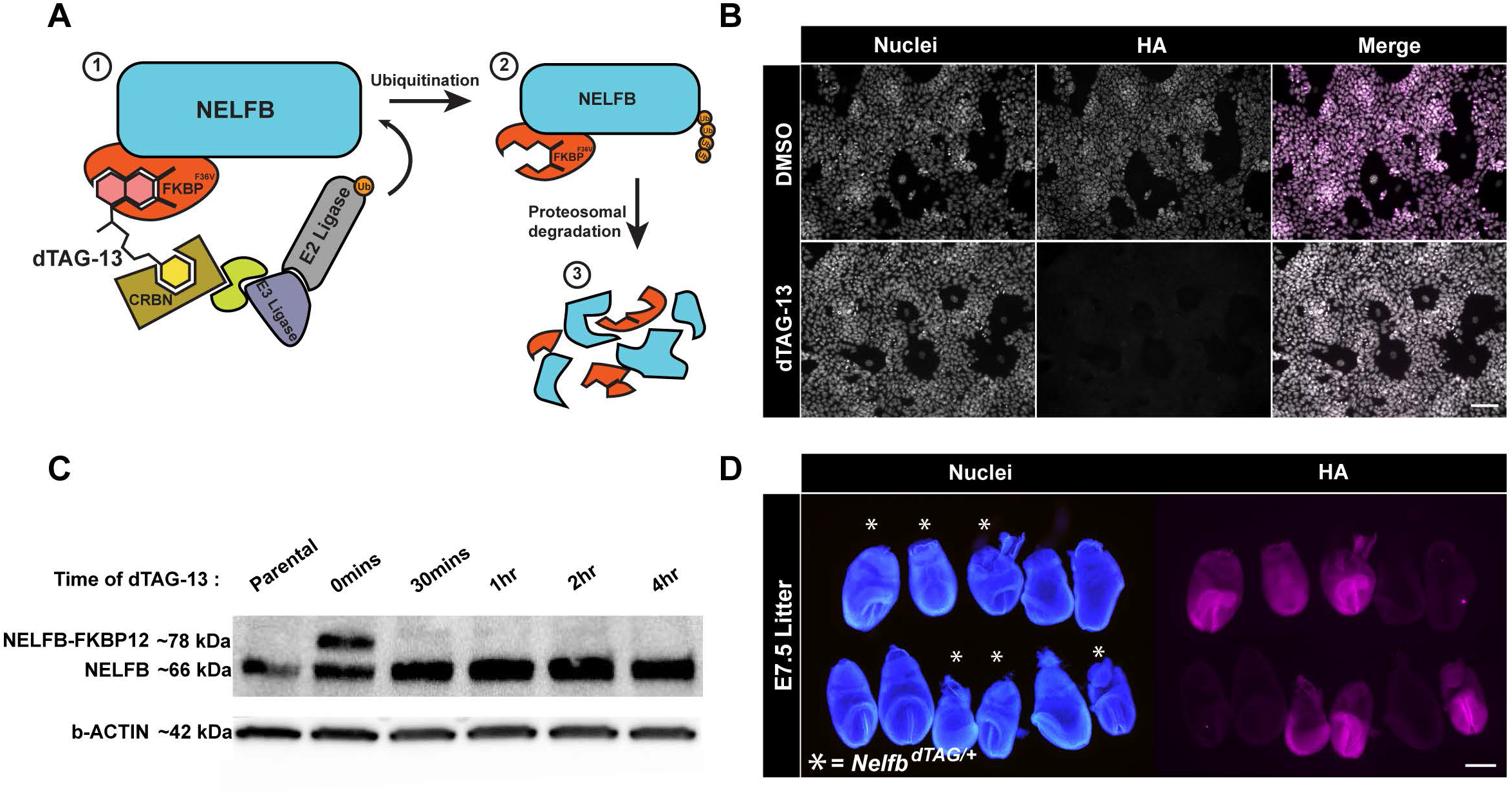
Generation of *Nelfb^dTAG^ mouse mod*el to study the dTAG system *in vivo*. (A) Schematic illustration of the proteasomal degradation in the dTAG system. The numbers represent the sequence of events from recognition of tag to degradation. (B) Immunofluorescence of targeted mESCs showing nuclear localized HA signal corresponding to NELFB +/- 500nM dTAG-13 treatment for 30 mins. Nuclei labeled with Hoechst. Scale bar, 70μm. (C) Western blot of targeted mES cells’ whole cell lysates +/- dTAG-13 treatment for indicated time intervals. Anti-NELFB antibody was used. 20ug of protein loaded/lane. (D) Immunofluorescence of a single E7.5 litter from a heterozygous *NelfbdTAG/+* male and wild-type *Nelfb^+/+^* female showing germline transmission of the targeted allele. Nuclei labeled with Hoechst. Scale bar, 500μm.

To test whether NELFB-FKBP^F36V^ could be degraded, we treated cells with 500nM dTAG-13. dTAG-13 is a heterobifunctional small molecule that binds to the FKBP^F36V^ tag and CRBN, resulting in rapid proteasomal degradation of the target protein (Figure 1A)(Nabet et al., 2018). As expected, NELFB-FKBP^F36V^ was rapidly degraded to undetectable levels when assayed via western blot or immunofluorescence (Figure 1B and 1C). Notably, full degradation was achieved within 30 minutes of adding dTAG-13, reflecting the temporal advantages of the dTAG system in mESCs.

To generate a mouse with the *Nelfb^dTAG^ allele, c*loned heterozygous mESCs containing the correctly targeted insertion were injected into 8-cell embryos to generate germline transmitting chimeras (Figure S1C)(Poueymirou et al., 2007). To test whether the edited allele is expressed in vivo, we performed immunofluorescence staining of the HA tag in E7.5 embryos resulting from heterozygous, *NelfbdTAG/+*, and wild-type, *Nelfb^+/+^*, crosses. These data showed that the *Nelfb^dTAG^ allele is* inherited at mendelian ratios and robustly expressed across heterozygous embryos, while HA staining is absent in wild-type embryos (Figure 1D). To further test the functionality of the tagged allele, heterozygous mice were crossed to generate homozygous *Nelfb^dTAG/dTAG^* mice. Heterozygous intercrosses resulted in viable and fertile homozygous animals that were indistinguishable from heterozygous and wild-type littermates, and were recovered at Mendelian ratios (Figure S1D). *Nelfb^dTAG/dTAG^* animals did not display any detectable phenotype and were able to generate normal sized litters (Figure S1E). Furthermore, complementing a previously generated null Nelfb allele with *Nelfb^dTAG^ resulted* in viable and healthy animals (Figure S1F)(Williams et al., 2015). Of note, Nelfb has been shown to be required for post-implantation mammalian embryonic development, suggesting that *Nelfb^dTAG^ is functi*onal (Amleh et al., 2009; Williams et al., 2015). These data support that we have generated a mouse model that widely expresses an endogenous NELFB-FKBP^F36V^ fusion protein. This enables us to evaluate the dTAG system in vivo and to glean insights into the role of NELFB in early development.

### The dTAG system is safe, efficient, and reversable in pre-implantation embryos

Mammalian pre-implantation development encompasses the time from fertilization to implantation of the embryo in the maternal uterine wall. During this period embryos grow independently from maternal tissues, and can be retrieved and routinely cultured ex utero (Piliszek et al., 2011). This flexibility has made pre-implantation embryos an excellent system to study a variety of biological processes in mice (Saiz et al., 2015). Before testing the efficiency of the dTAG system in pre-implantation embryos, we wanted to test whether dTAG-13 is safe for culturing mouse embryos from the zygote until the late blastocyst stages in vitro. Wild-type zygotes were cultured ex utero for 96 hours in standard medium (KSOM) with or without 500nM of dTAG-13 (Figure 2A)(Behringer et al., 2014). Based on our mESC experiments and previous reports, we expected that this concentration of dTAG-13 would be sufficient to activate degradation in pre-implantation embryos (Nabet et al., 2018, 2020). We found that dTAG-13 had no effect on the developmental potential of zygotes which developed to the late blastocyst stage (Figure S2A and S2B). To examine whether the ex utero cultured blastocysts with or without dTAG-13 developed normally, they were fixed and immunostained for markers of distinct cell fates. Late stage blastocyst embryos normally contain three cell lineages: epiblast (marked by NANOG), primitive endoderm (marked by GATA6), and trophoblast (marked by CDX2) (Schrode et al., 2013). We found that blastocysts that were cultured in the presence of dTAG-13 contained all these lineages (Figure S2C). Additionally, we found that the total cell number, and the ratio of trophoblast to total cell number were preserved (Figure 2A). Taken together these data suggest that, at concentrations shown to promote degradation in other systems, dTAG-13 is safe for pre-implantation development.

**Figure 2.**
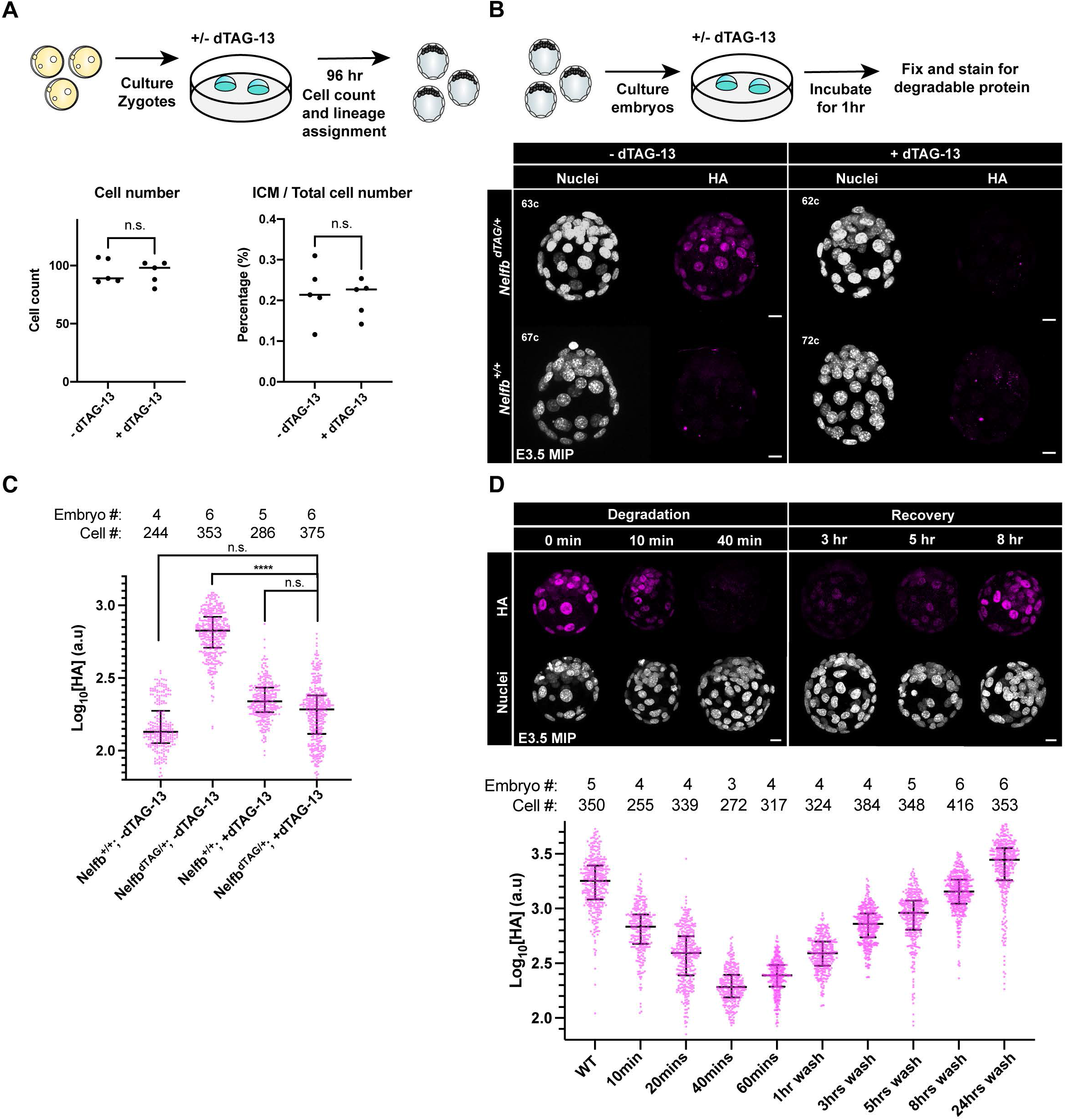
Establishing safety and efficiency of the dTAG system in pre-implantation stages. (A) (Top) Schematic representation of experimental design to test dTAG-13 safety in pre- implantation culture from zygote to blastocyst stages. (Bottom) Dot plots showing total cell counts (left) or percent ICM cells (right). Cell counts were determined following Hoechst staining and nuclei counting. ICM cells were identified via NANOG/GATA6 staining. Individual dots represent single embryos. Bars represent group means. (B) (Top) Schematic representation of experimental design to determine efficacy of tagged protein degradation. E3.5 blastocysts were cultured +/- 500nM dTAG-13 for 1hr prior to fixation and immunostaining. (Bottom) Immunofluorescence of HA to determine degradation of tagged protein. Nuclei labeled with Hoechst. Total cell count per embryo is shown in the top left corner. Scale bar, 10μm. (C) Quantification of mean HA signal intensity per nucleus. Wild type embryos used to show background fluorescence with staining. (D)(Top) E3.5 immunofluorescence of HA to determine degradation of tagged protein following several indicated incubation periods or following degradation and recovery in dTAG-13-free medium. Nuclei labeled with Hoechst. (Bottom) Quantification of mean HA signal intensity per nucleus. Scale bar, 10μm. For all experiments, maximum intensity projection (MIP) is shown in images. Plots show each data point with group mean and interquartile range. Student t-test was used to determine significance. Statistical significance is classified based on p-value as: n.s. > 0.05, * < 0.05, ** < 0.01, *** < 0.001, **** < 0.0001.

To test whether the dTAG system drives efficient degradation in pre-implantation embryos, we collected E3.5 blastocysts recovered from crosses of *NelfbdTAG/+* males with wild-type females. This crossing strategy provided both negative and positive controls within each litter for comparable quantitative analyses. Blastocysts were cultured for 1 hr in KSOM with or without dTAG-13. Embryos were then fixed and immunostained for HA to detect NELF-FKBP^F36V^. We found no detectable HA signal in treated *NelfbdTAG/+* embryos, suggesting highly efficient and rapid degradation (Figure 2B, 2C and S2D). Of note, degradation was observed across the entire embryo, and did not spare certain cells based on fate or position. Similar results were observed for E4.5 blastocysts (Figure S2F). The treatment had no acute effect on total cell number per embryo (Figure S2E). Performing the same experiment at finer temporal resolution, we noted that degradation was complete within 40 minutes of first exposure to dTAG-13 (Figure 2D). These results suggest that the dTAG system is efficient and can be used as an alternative strategy to conditional gene knockouts in pre-implantation embryos.

A key advantage of protein degradation approaches is their reversibility (Nabet et al., 2018). To test reversibility with the dTAG system in mouse pre-implantation embryos, E2.5 or E3.5 embryos were cultured in KSOM in the presence of dTAG-13 for 1 hr, followed by washing and culturing without dTAG-13 for various periods of time. Embryos were then fixed and immunostained for analysis. We found that the dTAG system is reversable within a few hours, with about 50% expression of degradable protein recovered in 5-8 hours (Figure 2D). While reversibility is slower than induction, and depends on additional factors such as levels of active gene expression, it is still rapid and outperforms other RNA-based systems such as RNAi (Verma et al., 2020). We speculate that the slower reversibility is in part due to potent binding of dTAG-13 to CRBN, thus requiring turnover of the protein to reverse the system. Taken together, these data show that the dTAG system is safe, rapid, efficient, and reversable in mouse pre-implantation embryos.

### The dTAG system is safe, efficient, and reversable in post-implantation embryos

Post-implantation development commences once the embryo implants into the maternal wall. In mice, this period starts around day E5.0. Early post-implantation development is a crucial time with intricate processes taking place such as pluripotency exit, onset of gastrulation and the initiation of organogenesis which occur within a 48 hour period in mice (Bardot and Hadjantonakis, 2020). Inducible and tissue-specific knockout systems, for example those driven by the lineage-specific and/or inducible forms of the Cre recombinase, have enabled discoveries during these embryonic stages. However, induction in such systems can be mosaic, inducing recombination in only a proportion of cells, especially before E8.5, and can be highly teratogenic resulting in abortion and mothers’ death (Pimeisl et al., 2013; Savery et al., 2020; Ved et al., 2019). These challenges make it difficult to generate inducible knockouts and study the consequences of acute protein loss during the early post-implantation period. Therefore, we aimed to test the safety and efficacy of the dTAG system at early post-implantation stages.

To test the safety of the dTAG system for post-implantation development, pregnant wild-type females received a single intraperitoneal (IP) injection of vehicle or dTAG-13 (35mg kg^-1^) at 6.5dpc, 8.5dpc, or 10.5dpc and were then allowed to go to term to assess any developmental defects resulting from treatments. We found that all mice delivered healthy litters in numbers comparable to a control group (Figure S3A). This experiment was repeated with dTAGv-1, a small molecule that activates the dTAG system via the VHL degradation path and reported to have improved in vivo pharmacokinetics than dTAG-13 (Nabet et al., 2020). Similarly, we found that dTAGv-1 had no effect on the health of pregnant females or size of their litters (Figure 3A). Thus, when administered to pregnant females both dTAG-13 and dTAGv-1 are developmentally neutral and safe to use during post-implantation stages of development.

**Figure 3.**
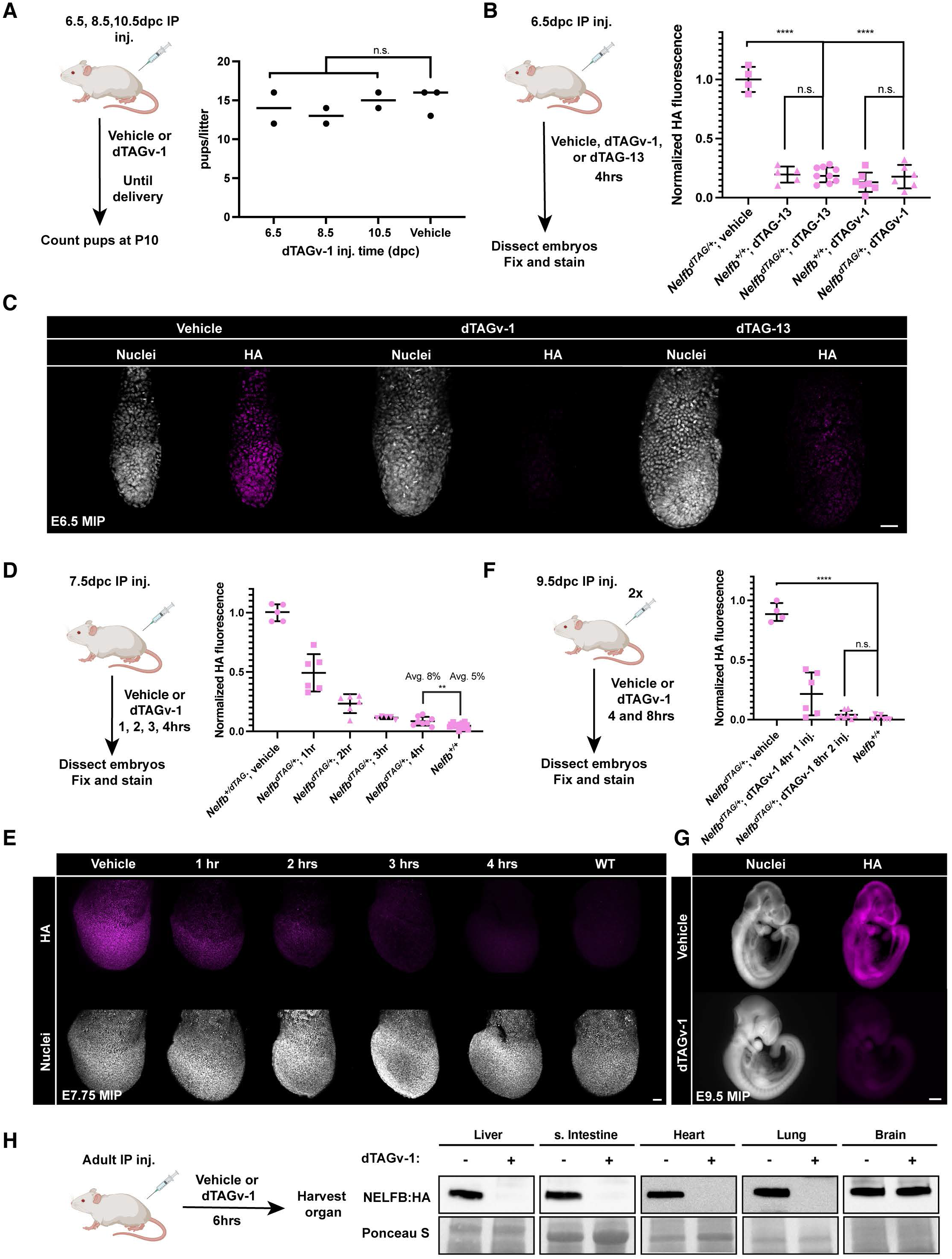
Establishing safety and efficiency of the dTAG system in post-implantation stages. (A) (Left) Schematic of testing dTAGv-1 safety in post-implantation development in vivo. Wild type pregnant females were given IP injection of 35mg kg^-1^ dTAGv-1 at 6.5, 8.5, 10.5 days post conception (dpc). (Right) Resulting healthy pups at term were counted. (B) Quantification of HA mean signal intensity in E6.5 embryos 4hrs after heterozygous *NelfbdTAG/+* pregnant females received dTAGv-1 or dTAG-13 injection. Each dot represents an embryo. HA signal was normalized to Hoechst, then vehicle injected litters. (C) Immunofluorescence images of E6.5 embryos retrieved 4hrs after pregnant females received IP injection of dTAGv-1 or dTAG-13. Nuclei are labeled by Hoechst. Scale bars, 50μm. (D)Quantification of HA mean signal intensity in E7.5 embryos 1,2,3, or 4hrs after heterozygous *NelfbdTAG/+* pregnant females received dTAGv-1 injection. Each dot represents an embryo. HA signal was normalized to Hoechst, then vehicle injected litters. (E) Immunofluorescence images of E7.5 embryos retrieved 1,2,3 and 4hrs after pregnant females received IP injection of dTAGv-1. Nuclei are labeled by Hoechst. Scale bars, 50μm. (F) Quantification of HA mean signal intensity in E9.5 embryos 4 and 8hrs after heterozygous *NelfbdTAG/+* pregnant females received one dTAGv-1 injection, or two separated by 2hrs and followed for 6hrs after the second for a total 8hrs from first injection. Each dot represents an embryo. HA signal was normalized to Hoechst, then vehicle injected litters. (G) Immunofluorescence images of E9.5 embryos retrieved 8hrs after pregnant females received two IP injections of dTAGv-1. Nuclei are labeled by Hoechst. Scale bars, 500μm. (H)Western blot of HA signal levels in several organs’ lysates 6hrs after IP injection of dTAGv-1. For all experiments, maximum intensity projection (MIP) is shown in images. Plots show each data point with group mean and interquartile range. Student t-test was used to determine significance. Statistical significance is classified based on p-value as: n.s. > 0.05, * < 0.05, ** < 0.01, *** < 0.001, **** < 0.0001.

Next, we wanted to test the efficiency of the dTAG system in vivo in post-implantation embryos. To do so, we delivered dTAG-13 or dTAGv-1 via IP injection to 6.5dpc pregnant females. After 4 hours, embryos were recovered, fixed, and stained. We found that a single injection of dTAG-13 or dTAGv-1 achieved complete and global degradation in E6.5 embryos (Figure 3B and 3C). To determine the dynamics of degradation, we performed the same experiment with females at 7.5dpc and collected embryos at 1, 2, 3, and 4 hours post injection. We observed >90% and >95% degradation was achieved within 3 and 4 hours respectively (Figure 3D and 3E). Similar results were obtained at mid-gestation stages when females at 9.5dpc were subject to a two-injection regime separated by 2 hours and followed for 6 hours post the second injection (Figure 3F and 3G). To determine whether the system is reversible in vivo, we injected animals with dTAG-13 or dTAGv-1 followed by a 24 hour chase before analysis. We found that expression was largely recovered at ∼70% with dTAG-13 or dTAGv-1 (Figure S3B and S3C). These results are in agreement with known pharmacokinetics of dTAG small molecules (Nabet et al., 2020).

Lastly, we sought to assess the functionality of the system in adult tissues. We found that a single injection of dTAGv-1 was sufficient to drive complete protein degradation within 6 hours in the liver, small intestine, lungs, and heart (Figure 3H). Notably, protein degradation was not detected in the brain, which is most likely protected from dTAG small molecules by the blood brain barrier. Taken together, these data demonstrate the dTAG system to be efficient and reversible during post-implantation mouse embryo development in vivo, and in adult animals.

### Acute degradation of NELFB reveals an essential role in pre-implantation development

Given the adaptability of the dTAG system to mammalian development and its unique, acutely inducible and reversable onset, we wanted to apply this system to a biological question that exploits these advantages. NELFB is an essential subunit of the negative elongation factor (NELF) complex (Yamaguchi et al., 1999). NELF has been shown to play a role in stabilizing transcriptionally engaged RNA Pol II at the promoter-proximal pausing position (Kwak and Lis, 2013; Yamaguchi et al., 1999). Upon phosphorylation by CDK9, NELF is released from the paused polymerase, enabling binding of pro-elongation factors (Adelman and Lis, 2012; Vos et al., 2018c, 2018b). Pol II pausing has been shown to play an essential role in the regulation of gene expression across a variety of biological contexts (Core and Adelman, 2019). In development, Nelfb knockout in mice results in embryonic lethality at early post-implantation in mice (Amleh et al., 2009). Furthermore, classical knockout strategies of essential genes cannot be used to examine whether a gene plays a role prior to the 4-8-cell stage in pre-implantation development, due to maternally deposited transcripts which can mask any potential phenotypes (Tadros and Lipshitz, 2009). A recent study has shown that Pol II pausing is uniquely prevalent in mouse embryos at the transition from zygote to 2-cell stages, coinciding with zygotic genome activation, or ZGA (Liu et al., 2020). Given that NELF subunits can be transcriptionally detected throughout pre-implantation mammalian development, we hypothesized that despite being dispensable for pre-implantation development after the 2-cell stage, NELF could play a critical role in ZGA.

To test this hypothesis, we first wanted to assess whether NELFB can be detected at the protein level in zygotes and 2-cell stage embryos. Previous analyses suggest that the Nelfb transcript is detected at low levels, but immunofluorescence studies have been inconclusive (Hu et al., 2020; Liu et al., 2020). We crossed homozygous *Nelfb^dTAG/dTAG^ male*s and females and collected zygotes at E0.5. Immunofluorescent staining for the HA tag revealed a signal localized to both maternal and paternal pro-nuclei (Figure 4B). Upon culture for 1 hour in dTAG, the signal was completely lost, suggesting degradation and activity of the dTAG system in zygotes (Figure 4A and 4B). Similar results were observed at the 2-cell stage (Figure S4A and S4B). The HA/NELFB staining at the zygote stage suggests that it is maternally deposited and expressed prior to ZGA. Because these embryos are the result of crossing homozygous mice, both maternal and zygotic transcripts are tagged, thus degradation ensures depletion of all protein present, regardless of the source.

**Figure 4.**
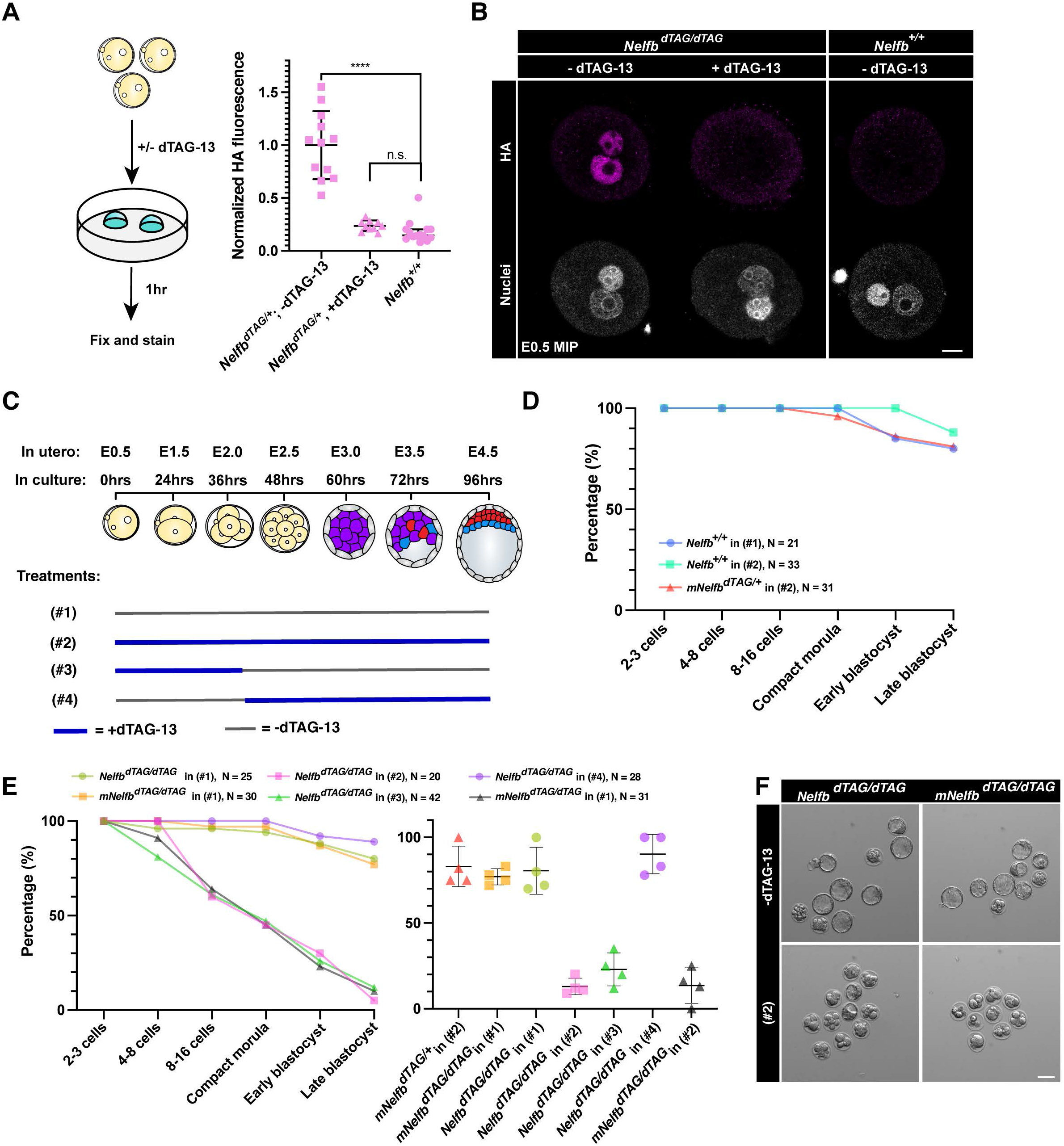
NELFB is required for pre-implantation development during zygote-2-cell stage. (A) (Left) Schematic of testing the dTAG system in zygotes. Homozygous *Nelfb^dTAG/dTAG^ embr*yos were collected at E0.5 and cultured with or without 500nM dTAG-13 for 1hr. (Right) Quantified mean HA signal in zygotes +/- 1 hr dTAG-13. Each dot represents a pro-nuclei. HA signal is normalized to non-treated embryos. (B) Immunofluorescence images of zygotes retrieved after 1hr culture +/- dTAG-13. Pro nuclei were labeled with Hoechst. Scale bars, 20μm. (C) Schematic of culture conditions for following panels, showing when dTAG-13 is added and removed for each condition in zygote to blastocyst culture. The treatment numbers: (#1): no dTAG-13, (#2) constant dTAG-13, (#3) 36hrs dTAG-13 from zygote to 4-cell stage followed by washing, (#4) 60hrs dTAG-13 from 4-cell stage to late blastocyst stage. (D) Average number of embryos reaching the indicated developmental stage per group/treatment. Control conditions are shown. Figure legends show genotype, treatment regiment, and total number of cultured embryos. (E) (Left) Average number of embryos reaching the indicated developmental stage per group/treatment. Control conditions are shown. Figure legends show genotype, treatment regiment, and total number of cultured embryos. Testing conditions are shown with littermate controls in each experiment. (Right) plot shows the average number of embryos successfully reaching late blastocyst stage in each separate litter per condition. (F) Representative images of cultured embryos at hour 96. Scale bars, 50μm. For all experiments, maximum intensity projection (MIP) is shown in images. Plots show each data point with group mean and interquartile range. Student t-test was used to determine significance. Statistical significance is classified based on p-value as: n.s. > 0.05, * < 0.05, ** < 0.01, *** < 0.001, **** < 0.0001.

To test whether NELFB plays a role in pre-implantation embryo development, we leveraged ex utero embryo culture between the zygote and blastocyst stages. To rigorously determine the safety of the dTAG system, we cultured wild-type and *NelfbdTAG/+* embryos (derived from *NelfbdTAG/+* mothers) in dTAG-13 for 96 hours, from the zygote to the late blastocyst stages. Both groups will be exposed to dTAG-13, but the *NelfbdTAG/+* embryos will also have the degradation system engaged since they harbor one *Nelfb^dTAG^ allele. U*nder these conditions embryos developed to late-stage blastocysts at a comparable rate (∼80%) to wild-type embryos in the absence of dTAG-13 (Figure 4D and S4C). By contrast, culturing homozygous embryos in dTAG resulted in severe developmental arrest of ∼80% of embryos at stages prior to the blastocyst stage (Figure 4C, 4E and 4F). Of note, about 50% were arrested at early cleavage stages, suggesting an early requirement for NELFB in pre-implantation development. To test whether this requirement is specific to the zygote/2-cell stages, we cultured homozygous *Nelfb^dTAG/dTAG^ embr*yos in dTAG from the zygote until the 4-cell stage, or from 4-cell until late blastocyst stage (Figure 4C). An early treatment recapitulated the developmental phenotype of dTAG treatment when present during the entire culture period, while the late treatment had no effect (Figure 4E and S4D). Furthermore, culturing maternal homozygous *Nelfb^dTAG/dTAG^; zyg*otic heterozygous *NelfbdTAG/+* embryos (resulting from crossing *Nelfb^dTAG/dTAG^ fema*les with *Nelfb^+/+^* males) in dTAG recapitulated the *Nelfb^dTAG/dTAG^ homo*zygous embryo defect (Figure 4E and 4F). These experiments suggest that NELFB, and likely maternally provided NELFB, is required for pre-implantation development specifically during the transition from zygote to 2-cell stage, and is dispensable thereafter.

### NELFB degradation at the zygote-2-cell stages compromises ZGA

Based on our observation that NELFB is required specifically at the zygote-2-cell stage, we hypothesized that NELFB degradation alters ZGA in mouse embryos. To test this hypothesis, *Nelfb^dTAG/dTAG^ litt*ermate zygotes were collected and cultured for 30 hours, corresponding to late 2-cell stage, in the presence or absence of dTAG-13. Embryos were then collected from each condition in two replicates, each replicate containing three embryos, for low-input SMART-seq to analyze gene expression (Figure 5A). Samples clustered based on treatment (Figure S5A and S5B). Overall, the expression patterns of 2610 genes were significantly changed (p. adj. < 0.05), with ∼53% downregulated (Figure 5B and S5C). Among the differentially expressed genes were ones specific to the 2-cell state and ZGA-associated factors, such as Zscan4e, Zscan4d, and Dppa4 (Figure 5B)(Hu et al., 2020).

**Figure 5.**
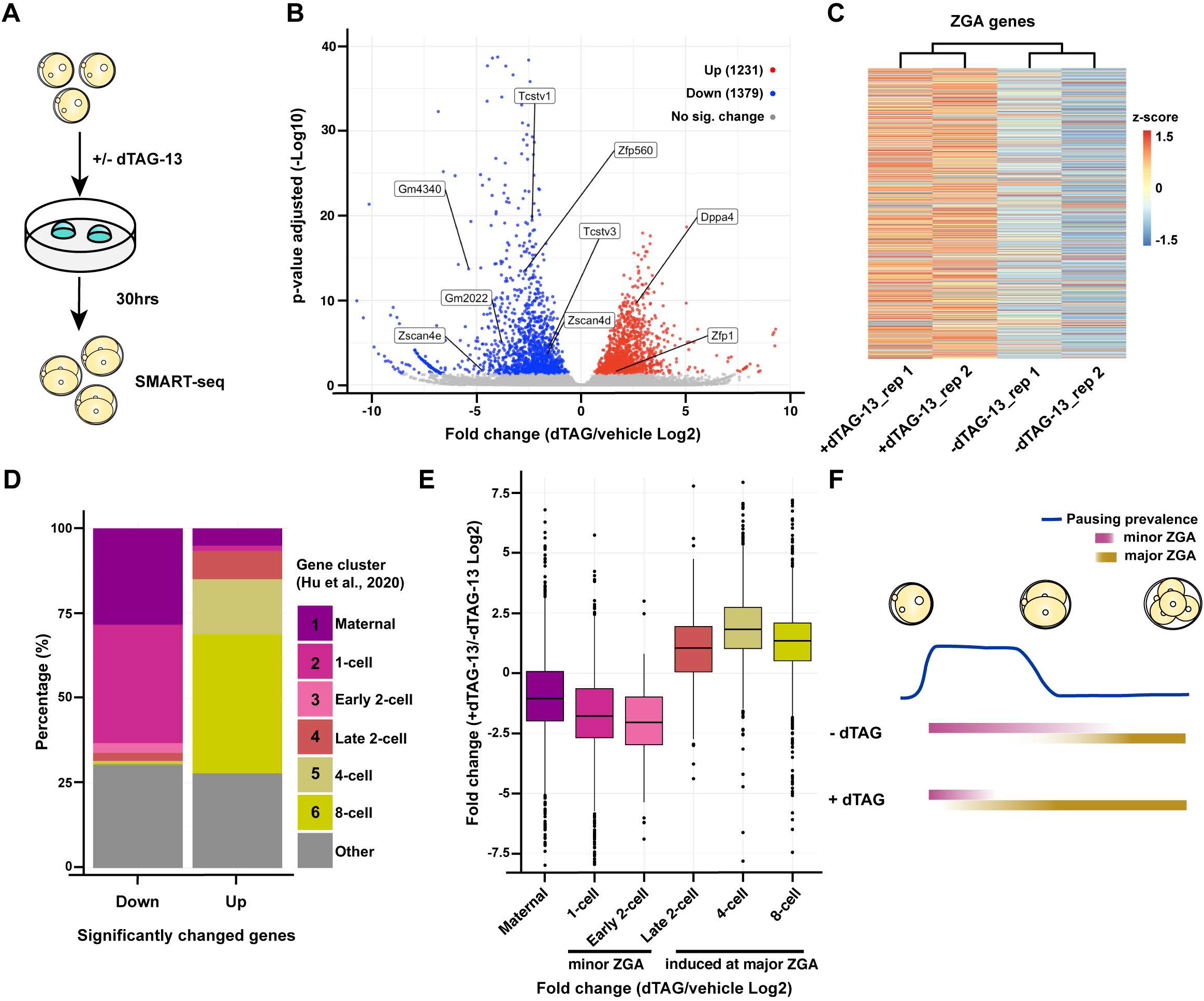
NELFB facilitates ZGA in mouse development. (A) Schematic of experiment. *Nelfb^dTAG/dTAG^ zygo*tes were cultured for 30 hr +/- dTAG-13 to late 2-cell stage, then collected for sequencing using SMART-seq. (B) Volcano plot of differentially expressed genes determined by DEseq2. P. adj of 0.05 was used as a cutoff. (C) Heatmap showing z-score of major ZGA genes +/- dTAG-13. 3481 genes are shown representing cluster one from Abe et al., 2018. (D) Identifying the clusters of up- and downregulated genes using clusters from Hu et al., 2020. (E) Boxplot showing the overall change of expression levels +/- dTAG-13 in each cluster from Hu et al., 2020. (F) Working model to explain the results. NELF may serve to stabilize RNA Pol II pausing prior to major ZGA, resulting in major ZGA attenuation.

To identify the classes of differentially expressed genes, we used gene clusters present in zygote-to-8-cell stage embryos identified in two previous studies (Abe et al., 2018; Hu et al., 2020). A broad analysis of ∼3000 ZGA-associated genes revealed their overall upregulation in dTAG-13 treated samples (Figure 5C). To further classify the specific identity of upregulated and downregulated genes, we queried the stage-specific clusters (Hu et al. 2020) to which these genes belonged to. Over 50% of downregulated transcripts belonged to genes that peak in the zygote-to-early-2-cell state (Figure 5D). By contrast, over 50% of upregulated genes belonged to major ZGA genes that are initially expressed in late 2-cell stage and increase expression until the 4- and 8- cell stages (Figure 5D). When assessing the overall gene expression levels in each cluster, we find that there is a general increase in clusters representing major ZGA and a decrease in clusters representing minor ZGA (Figure 5E). We obtained similar results when using clusters identified in a different study (Abe et al. 2018, Figure S5D and S5E). Taken together, these results suggest that stage-specific NELFB degradation alters ZGA gene expression, which may result in a compromised minor ZGA and premature major ZGA. These data corroborate previous studies showing that pausing peaks at the zygote-to-early-2-cell stage and decreases at the late 2-cell stage, suggesting that NELFB plays a role in pausing at this stage (Liu et al., 2020).

## DISCUSSION

Targeted protein degradation has recently emerged as a method to study protein function in cultured cells (Wu et al., 2020). Here, we successfully adapted one of the promising protein degradation systems, the dTAG system, to the mouse model (Nabet et al., 2018, 2020). Specifically, we show that the dTAG system is efficient and safe across mouse development when administered to embryos in ex utero culture or via introduction to pregnant females.

One of the main advantages of protein degradation is the onset of effect which typically occurs within minutes to hours (Clift et al., 2017; Nabet et al., 2018; Nishimura et al., 2009). This is a major advantage over other RNA/DNA targeting techniques. Our work demonstrates that these rapid dynamics can be achieved in ex utero cultured pre-implantation embryos, in vivo in post-implantation embryos, and even adult organs with a single IP injection. Notably, near complete degradation of our targeted protein was consistently observed across all these stages, suggesting that the dTAG system is a viable and attractive approach to study acute protein ablation in an inducible manner. Furthermore, the modular nature of the dTAG system enables engaging different ubiquitination pathways with different small molecules which expands the potential tissues and cell lines where they can function (currently CRBN and VHL can be engaged with dTAG-13 and dTAGv-1 respectively)(Nabet et al., 2018, 2020).

The dTAG system can overcome several challenges that are unique to studying protein function in mouse development. One example being transcripts that are maternally deposited prior to zygotic genome activation. Using the dTAG system, one can circumvent the need to use a floxed allele and an oocyte-specific Cre recombinase driver to study essential genes in pre-implantation development. Since tagged alleles can be normally expressed and functional in the absence of dTAG small molecules, it is possible to retrieve 100% of embryos from homozygous crossings that can be induced to be maternal and zygotic deficient of a certain protein. This results in significantly reduced time and resources to obtain maternal/zygotic depleted embryos, and simultaneously a higher yield. Importantly, the rapid inducible and reversable nature of this system enables fine control to define the exact window of necessity of a protein during development and the potential to perform successive depletion-and-recovery cycles. Another challenge that can be overcome with the dTAG system is the inducible depletion of proteins at early post-implantation stages (E5.5 – E7.5). Using CreER systems to study gene function at these stages can be limited by the mosaicism and teratogenicity of tamoxifen (Pimeisl et al., 2013; Ved et al., 2019).

By leveraging the advantages of the dTAG system we gleaned new insights regards the role of NELFB and RNA Pol II pausing during pre-implantation development. Nelfb null embryos have been reported to exhibit an early post-implantation lethality, but a role for NELFB in pre-implantation development had not been documented (Amleh et al., 2009). By leveraging the temporal control afforded by the dTAG system with simple mouse genetics and one allele, we uncovered a key role for maternal Nelfb in controlling the minor and major ZGA. These observations are in agreement with the reported critical role of Nelfa in the Drosophila maternal-to-zygotic (MZT) transition (Wang et al., 2010). Additionally, our data suggest that NELFB regulates expression of minor ZGA genes, and is dispensable for major ZGA, in agreement with the recently reported pausing enrichment prior to major ZGA (Liu et al., 2020). These observations open the door to further studies aimed at determining the molecular mechanisms driving this divergent control of distinct gene sets.

Taken together, our data demonstrate that the dTAG system offers unique advantages to other existing perturbation tools and can be applied efficiently in vivo across mammalian development.

### LIMITATIONS

We present here compelling evidence that the dTAG system is highly efficient and functional in early mammalian development. However, there are few limitations and considerations that should be highlighted. First, as is the case with any other system, efficiency and dynamics of the dTAG system are partially target dependent (Nabet et al., 2018, 2020). Thus, it is necessary to validate degradation of the target protein in cell models before generating mice. Importantly, a fusion protein-tag is required for the system, making it possible that the tag may misfold or impact the protein’s function. Thus, it is necessary to validate the function of the tagged protein.

One other in vivo-specific limitation is the pharmacokinetics of dTAGv-1 and dTAG-13. Our data suggests that to maintain full degradation of the target protein in vivo, injections are required every ∼12 hours. Thus, while the system is ideal for short-term studies, it can be cumbersome for long-term studies. This can be potentially circumvented by continuous perfusion systems or development of longer acting in vivo dTAG small molecules.

## ACKNOWLEDGEMENTS

We thank MSK’s Mouse Genetics Core Facility for assistance in generating *Nelfb^dTAG^ mice and* the Integrated Genomics Operation and Bioinformatics Core Facility for assistance in sequencing and sequence data analysis. We are grateful to members of the Hadjantonakis lab for stimulating discussion and critical feedback. AA is supported by a MSTP training grant from the NIH (T32GM007739) awarded to the Weill Cornell/Rockefeller/Sloan Kettering Tri-Institutional MD-PhD Program and NIH F30HD103398. Work in AKH’s lab is supported by the NIH (R01HD094868, R01DK127821, R01HD086478, and P30CA008748).

## AUTHOR CONTRIBUTIONS

Conceptualization, A.A. and A.K.H.; Methodology, A.A.; Investigation; A.A., Data Curation, A.A.; Writing – Original Draft, A.A.; Writing – Review & Editing, A.A. and A.K.H.; Funding Acquisition, A.A. and A.K.H.; Supervision, A.K.H.

## STAR METHODS

### Lead contact and materials availability

Request for reagents should be directed to and will be fulfilled by the lead contact, Anna-Katerina Hadjantonakis

### Experimental model details

#### Cell lines

C57BL/6-derived HK3i ES cell line was cultured on 0.1% gelatin (Millipore) coated tissue-culture grade plates in a humidified 37°C incubator with 5% CO_2_ (Kiyonari et al., 2010). For routine culture, cells were grown in DMEM (Gibco), supplemented with 2mM L-glutamine (Gibco), 1x MEM non-essential amino acids (Gibco), 1mM sodium pyruvate (Gibco), 100U/ml penicillin and 100μg/ml streptomycin (Gibco), 0.1mM 2-mercaptoethanol (Gibco), 15% KnockOut^TM^ Serum Replacement (Gibco), and 1000U/ml of recombinant leukemia inhibitory factor (LIF). ESCs were maintained on a layer of Mitomycin C inactivated mouse embryonic fibroblasts (MEFs).

#### Plasmid generation

Two plasmids were generated for this study: (1) Cas9 vector to target the C-terminus of Nelfb gene. PX459 vector (addgene #62988) was digested using BbsI-HF (NEB) and single guide RNA targeting Nelfb was annealed (Ran et al., 2013), (2) Homology directed repair (HDR) vector containing the insert FKBP^F36V^ tag, 2x HA tag, self-cleaving P2A sequence, and Puromycin resistance, flanked by 1 kb Nelfb HDR sequences. The insert was obtained from pCRIS-PITCHv2-dTAG-Puro (addgene #91796)(Nabet et al., 2018). The plasmid backbone (pBluescript), Nelfb HDR sequences, and the insert were amplified using Q5 polymerase (NEB) and the plasmid was constructed using NEBuilder HiFi DNA assembly (NEB).

#### Genetic editing

To generate *NelfbdTAG/+* mESCs, 3 million cells were transfected with 10ug PX459- Nelfb_sgRNA and 10ug Nelfb_left- FKBP^F36V^- 2xHA- P2A-Puro-Nelfb_right. Cells were transfected using Lonza P3 Primary Cell 4D-Nucleofector^TM^ X 100ul cuvettes (Lonza).

Following transfection, cells were plated on a 10 cm dish (Falcon) coated with MEFs. 48 hrs post transfection, correctly targeted cells were selected for using 1.8 ug/ml puromycin (InvivoGen) for 48 hrs. Surviving cells were split 1000 cells/10 cm dish and maintained for 9 days under Puromycin selection. Surviving clones were picked under a stereomicroscope, expanded, and genotyped for the insert.

#### Mouse strains and husbandry

All animal work was approved by MSKCC Institutional Animal Care and Use Committee (IACUC). Animals were housed in a pathogen-free facility under a 12 hr light cycle. Mouse strains used in this study were *Nelfb^dTAG^ and wild-*type CD-1/ICR (Charles River). *Nelfb^dTAG^ mice were* generated by the Mouse Genetics Core at MSKCC. *NelfbdTAG/+* ESCs were injected into C57Bl/6 Albino host 8-cell morula. After culturing for 24 hrs in KSOM (Millipore), resulting blastocyst were transferred to pseudo-pregnant females. Only chimeras with ∼100% coat color contribution from injected cells were used and mated with CD-1/ICR mice (Chalres River). The resulting colony was outbred to CD-1 three times. Homozygous colony was then established using these mice.

### Method details

#### Cells dTAG treatment

dTAG-13 (Bio-Techne) was reconstituted in DMSO (Sigma) at 5mM. dTAG-13 was diluted in maintenance medium to 500nM and added to cells with medium changes for the specified amounts of time.

#### Embryo collection

For all experiments, embryos were obtained via natural mating of 6-12 weeks of age females with 7 – 16 weeks of age males. For preimplantation stages, embryos were recovered by flushing the uterine horns (E3.5 – E4.5), the fallopian tubes (E1.5 – E2.5), or dissecting cumulus cells from the fallopian tubes (E0.5). These dissections were carried out in flushing and holding medium (FHM, Millipore) as described (Behringer et al., 2014). To separate zygotes from cumulus cells, cumulus cells-covered embryos were incubated for ∼3 mins in 0.3mg/ml Hyaluronidase (Sigma) in FHM.

For post-implantation embryos (E6.5 – E9.5), the uterine horns were retrieved and cut into single decidual swellings in 5% Newborn Calf Serum in DMEM/F12 (Gibco). Embryos were dissected out by removing the uterus wall and decidual tissue. For E6.5-E7.5 embryos, the parietal endoderm was removed carefully with the ectoplacental cone. For E9.5 embryos, the amnion was carefully removed.

#### Pre-implantation embryo culture

Once recovered, embryos were washed 2 times and cultured in KSOM-AA without phenol red (Millipore). 5-15 Embryos were culture in a 60mm organ culture dish (Falcon) with 700ul KSOM-AA, and 4 ml H2O in the humidifying chamber. Dishes were incubated in a humidified 37°C incubator with 5% CO2 for up to 96 hrs. For degradation treatments, embryos were incubated with control (KSOM-AA and 0.01% DMSO) or 500nM dTAG-13 (Bio-Techne) in KSOM-AA for the duration of treatment.

#### Animal injections

For intraperitoneal (IP) injections, treatments were formulated in 10% Cremophore EL (Sigma) in sterile PBS+/+. dTAGv-1 (Bio-Techne)/dTAG-13 was initially reconstituted in DMSO at 1mg/25ul. Prior to injection, 25ul of dTAGv-1/13 or DMSO for vehicle was diluted in 475ul 10% Cremophore EL. Final injection volume was 0.5 ml of 5% DMSO, 10% Cremophore EL in PBS +/+ with or without 1mg dTAGv-1/13 to achieve ∼35mg/kg concentration/injection.

#### Immunofluorescence

For cultured ESCs, cells were plated on u-Slide 8 well (ibidi), washed with PBS+/+ and fixed in 4% PFA (electron microscopy sciences) in PBS+/+ for 10 min at room temperature. Fixed cells were washed two times with PBS+/+, one time with wash buffer; 0.1% Triton X- 100 (Sigma) in PBS+/+, then permeabilized in 0.5% Triton X-100 (Sigma) in PBS+/+ for 10 min. Cells were then blocked with 3% Donkey Serum (Sigma) and 1% BSA (Sigma) for 1 hr at room temperature. Cells were then incubated with mouse anti-HA (abcam, 1:500) in blocking buffer at 4°C over night. Cells were then washed three times in wash buffer, and incubated with donkey anti-mouse Alexa Fluor^TM^ 488 (Invitrogen, 1:500) for 1 hr at room temperature. Cells were then washed three time wish wash buffer, the last containing 5μg/ml Hoechst 33342 (Invitrogen), then imaged.

For E3.5-E4.5 pre-implantation embryos, the zona pellucida was removed by incubation in acid Tyrode’s solution (Sigma) at 37°C for 2 min. Embryos were subsequently washed briefly in PBS+/+ before fixation in 4% PFA for 10 mins at room temperature. Fixed embryos were washed in 0.1% Triton X-100 in PBS+/+ (PBX)for 5 min, permeabilized in 0.5% Triton X-100 (Sigma) in PBS+/+ for 5 min, washed again for 5 min in PBX, and blocked in 2% horse serum (Sigma) in PBS+/+ for 1 hr at room temperature. Embryos were incubated in primary antibodies diluted in blocking solution over night at 4°C. Embryos were then washed three times for 5 min each in PBX and blocked again for 1 hr at room temperature prior to incubation with secondary antibodies. Secondary antibodies diluted in blocking solution were applied for 1 hr at 4°C. Embryos were then washed twice for 5 min each in PBX and incubated with 5ug/ml Hoechst 33342 (Invitrogen) in PBS for 5 min or until mounting for imaging. The following primary antibodies were used: goat anti-GATA6 (R&D Systems, 1:100), mouse anti-CDX2 (BioGenex, 1:200), rabbit anti-NANOG (CosmoBio, 1:500), mouse anti-HA (abcam, 1:500), rabbit anti-HA (Cell Signaling, 1:200). Secondary Alexa Fluor-conjugated antibodies (Invitrogen) were used at a dilution of 1:500. DNA was visualized using Hoechst 33342.

For E0.5-E1.5 embryos, the staining was identical to E3.5-E4.5 embryos except for fixation for 50 min at room temperature in 4% PFA, permeabilization for 30 min at room temperature, and blocking solution is 1% BSA in PBS+/+.

For E6.5 and E 7.5, Embryos were washed briefly in PBS+/+ before fixation in 4% PFA for 20 mins at room temperature. Fixed embryos were washed in 0.1% Triton X-100 in PBS+/+ (PBX) for 5 min, permeabilized in 0.5% Triton X-100 (Sigma) in PBS+/+ for 20 min, washed again for 5 min in PBX, and blocked in 3% horse serum (Sigma) in PBX for 1 hr at room temperature. Embryos were incubated in primary antibodies diluted in blocking solution over night at 4°C. Embryos were then washed three times for 10 min each in PBX and blocked again for 1 hr at room temperature prior to incubation with secondary antibodies. Secondary antibodies diluted in blocking solution were applied over night at 4°C. Embryos were then washed three times for 5 min each in PBX and incubated with 5ug/ml Hoechst 33342 (Invitrogen) in PBX for 1 hr or until mounting for imaging.

For E9.5, embryos were fixed in 4% PFA for 40 mins at room temperature. Fixed embryos were washed in 0.1% Triton X-100 in PBS+/+ (PBX) for 15 min, permeabilized in 0.5% Triton X-100 (Sigma) in PBS+/+ for 1 hr, then blocked in 5% horse serum (Sigma) in PBX for 2 hr at room temperature. Embryos were incubated in primary antibodies diluted in blocking solution over night at 4°C. Embryos were then washed in PBX three times for 30 mins each and blocked again for 2 hr prior to incubation in secondary antibodies and Hoechst over night at 4°C. Embryos were subsequently washed three times for 30 mins each in PBX. Embryos were mounted in Ce3D clearing solution 4 hr before imaging (Li et al., 2019).

#### Image data acquisition from fixed samples

Fixed immunostained samples were imaged on a Zeiss LSM880 laser scanning confocal microscope. Pre-implantation embryos were mounted in microdroplets of 5μg/ml Hoechst 33342 in PBS+/+ on glass-bottomed dished (MatTek) coated with mineral oil (Sigma). Embryos were imaged along the entire z-axis with 1μm step using an oil-immersion Zeiss EC Plan-Neofluar 40x/NA 1.3 with a 0.17mm working distance. For post-implantation embryos, a similar setup was used but with an air Plan-Apochromat 20x/NA 0.75 objective.

Low magnification pooled embryo images and E9.5 full embryos were imaged on a Zeiss Axio Zoom.V16 using a PlanNeoFluar Z 1x/0.25 FWD 56mm objective. Pseudo-stacking (Zeiss ZEN Blue) was used for E9.5 images.

#### Western blotting

For cells, 350ul of lysis buffer; 1x Cell Lysis Buffer (Cell Signaling) with 1mM PMSF (Cell Signaling) and cOmplete^TM^ Ultra protease inhibitor (Sigma), was added to a 90% confluent 6-well dish (Falcone) after washing with PBS-/-. Cells were incubated with lysis buffer for 5 min on ice, then scraped and collected. Samples were sonicated for 15 seconds to complete lysis at, then spun down at 12,000x g for 10 min at 4°C. The supernatant was collected, and protein concentration measured using Pierce^TM^ BCA Protein Assay Kit (Thermo). 20ug of protein was mixed with Blue Loading Buffer (Cell Signaling) and 40mM DTT (Cell signaling). Samples were boiled at 95°C for 5 min for denaturation. For tissue samples, similar protocol was followed, except that 100mg of tissue was homogenized in 1ml lysis using a Dounce homogenizer for 30 mins on ice.

Samples were run on a BioRad PROTEAN system and transferred using Trans-Blot Semi-Dry Transfer Cell (BioRad) to a nitrocellulose membrane (Cell Signaling) following manufacturer’s instructions and reagents. Membranes were then washed briefly with ddH2O and stained with Ponceau S (Sigma) for 1 min to check for transfer quality, and as a loading control. Membranes were then washed three times with TBST; 0.1% Tween 20 (Fisher) in TBS. Membranes were blocked with 4% BSA in TBST for 1 hr at room temperature and subsequently incubated with primary antibodies diluted in blocking buffer at 4°C over night. They were then washed three times with TBST, then incubated with secondary antibodies in blocking buffer for 1 hr. Washed three times with TBST, incubated with ECL reagent SignalFire^TM^ for 1-2 min and imaged using a ChemiDoc (BioRad). The following antibodies were used: rabbit anti-HA (Cell Signaling, 1:1000), mouse anti-b-Actin (Cell Signaling, 1:3000), anti-rabbit IgG, HRP-linked (Cell signaling, 1:2000), anti-mouse IgG, HRP-linked (Cell signaling, 1:2000).

#### RNA transcriptomics

Sequencing was done by the MSKCC Integrated Genomics Operations (IGO). SMART-seq v4 Ultra Low Input pipeline was used (Clonetech/Takara). Three 2-cell stage embryos were added to 5ul of 10x Lysis Buffer (Takara) and RNase inhibitor (Takara) per manufacturer’s instructions. Samples were immediately frozen on dry ice and stored at -80 until cDNA amplification, library prep, and sequencing using NovaSeq 6000.

### Quantification and statistical analysis

#### Image processing and quantification

For Pre-implantation embryos, semi-automated 3D nuclear segmentation for cell counting and quantification of fluorescence intensity was carried out using MINS, a MATLAB-based algorithm (http://katlab-tools.org/) (Lou et al., 2014). The same imaging parameters were used for all experiments consisting of the same primary and secondary antibody combinations to minimize quantitative variance due to image acquisition. The MINS output was checked for over- or under-segmentation and tables were corrected manually using Image J (NIH, https://imagej.nih.gov/ij/). Under-segmented nuclei (two or more nuclei detected as one, or nuclei that were not detected) were assigned fluorescence intensity values that were directly measured using ImageJ (NIH). To correct fluorescence decay along the Z-axis, we used a linear regression method to calculate the global average of the regression coefficients in the HA channel (Saiz et al., 2016).This slope was then used to adjust the logarithm values of HA fluorescence intensity for each nucleus. Trophectoderm (TE) vs. inner cell mass (ICM) cell assignment was achieved by a threshold for CDX2 which is present exclusively in TE. To avoid batch variability, directly compared embryos were stained and imaged in the same session.

For post-implantation embryos, fluorescence intensities were acquired manually per embryo by measuring HA and Hoechst in ∼10 nuclei. HA signal was normalized to Hoechst per embryo, and all measurements were compared only to similar age embryos that were stained and imaged in the same session, except for E7.5 time series which was done over two sessions due to embryos number. All embryos were stained, imaged, and processed using the same parameters, and intensity quantifications were measured at the same Z-depth across all embryos to omit the need for Z- fluorescence decay normalization.

#### Statistical analysis

All statistical tests of immunofluorescence data were carried out in PRISM 9 (GraphPad). Statical significance was established using a two-tailed student t-test with p-value threshold of 0.05. The p-value range for each experiment is indicated in the figure legend.

For sequencing data, analysis of differentially expressed genes was done in R using the DEseq2 method (Love et al., 2014).

#### Data and code availability

Source data for all figures is available in the supplements, Table S2. RNA sequencing data from this work is available in the Gene Expression Omnibus under the accession number GSE185099.

### Key resources table

| REAGENT or RESOURCE | SOURCE | IDENTIFIER |
| --- | --- | --- |
| Antibodies |  |  |
| Rabbit anti-NANOG | Reprocell | Cat# RCAB001P, RRID: AB_1962694 |
| Goat anti-GATA6 | R&D Systems | Cat# AF1700, RRID: AB_2108901 |
| Mouse anti-CDX2 | BioGenex | Cat# MU-392AUC, RRID: AB_2650531 |
| Rabbit anti-COBRA1/NELFB | Invitrogen | Cat# PA5-54169, RRID: AB_2639998 |
| Mouse anti-HA | Abcam | Cat# ab130275, RRID: AB_11156884 |
| Rabbit anti-HA | Cell Signaling | Cat# 3724, RRID: AB_1549585 |
| Bacterial and Virus Strains |  |  |
| E. coli DH5a | NEB | C2987I |
| Chemicals, Peptides, and Recombinant Proteins |  |  |
| Hoechst 33342 | Invitrogen | Cat# H3570 |
| Stemolecule™ CHIR99021 | Fisher | Cat# NC9785126 |
| PD0325901 | Reprocell | Cat# 04-0006-10 |
| EmbryoMax KSOM (+AA w/o PhenolRed) | Sigma | Cat# MR-106-D |
| FHM | Millipore | Cat# MR-025-D |
| Proteinase K | Roche | Cat# 03115801001 |
| 16% PFA | Electron microscopy sciences | Cat# 15710 |
| Acid Tyrode's | Millipore | Cat# MR-004-D |
| Glycine | Sigma | Cat# G7403 |
| Triton X-100 | Sigma | Cat# X100 |
| Horse Serum | Sigma | Cat# H0146 |
| DMEM | Life technologies | Cat# 11995073 |
| NEAA | Life technologies | Cat# 11140-050 |
| Glutamine | Life technologies | Cat# 25030164 |
| Sodium Pyruvate | Life technologies | Cat# 11360070 |
| 2-mercaptoethanol | Life technologies | Cat# 21985023 |
| Fetal Calf Serum | VWR | Cat# 97068-085 |
| 0.05% Trypsin EDTA | Life technologies | Cat# 25200114 |
| Hyaluronidase | Millipore | Cat# H4272 |
| Gelatine | Millipore | Cat# 104070 |
| Penicillin/Streptomycin | Life technologies | Cat# 15140163 |
| Mitomycin C | Sigma | Cat# M4287 |
| Critical Commercial Assays |  |  |
| NEBuilder HiFi assembly kit | NEB | Cat# E5520S |
| Pierce™ BCA protein assay kit | Thermo | Cat# 23225 |
| Deposited Data |  |  |
| RNA-seq data | This paper | GSE185099 |
| All source data for figures | Table S2 | N/A |
| Experimental Models: Cell Lines |  |  |
| Mouse: Embryonic stem cell line HK3i | MSK Mouse Genetics Core Facility | (Kiyonari et al., 2010) |
| Experimental Models: Organisms/Strains |  |  |
| Mouse: CD1 | Charles River Laboratory | Cat# 022 |
| Oligonucleotides |  |  |
| Oligonucleotides in Table S1 | This paper | N/A |
| Software and Algorithms |  |  |
| Fiji/ImageJ | (Schindelin et al., 2012) | https://imagej.nih.gov/ij/ |
| R Studio 1.1.456/R 4.0.2 | RStudio, Inc. | https://rstudio.com/ |
| DESeq2 3.13 | Bioconductor (Love et al., 2014) | https://bioconductor.org/packages/release/bioc/html/DESeq2.html |
| ggplot2 3.3.5 | R package | https://ggplot2.tidyverse.org |
| MINS | (Lou et al., 2014) | http://katlab-tools.org/ |
| Prism 9 | GraphPad Software | https://www.graphpad.com/scientific-software/prism/ |
| ZEN | Carl Zeiss Microsystems | https://www.zeiss.com/microscopy/us/products/microscope-software/zen.html |

| Other |  |  |
| --- | --- | --- |
| Glass-bottom dish | MatTek | Cat# P35G-1.5-14-C |
| Nitrocellulose membrane | Cell Signaling | Cat #12369 |

## SUPPLEMENTAL TABLES AND FIGURE LEGENDS

**Table S1. List of oligonucleotides used in this study.**

**Table S2. Source data for all figures.**

**Figure S1.**
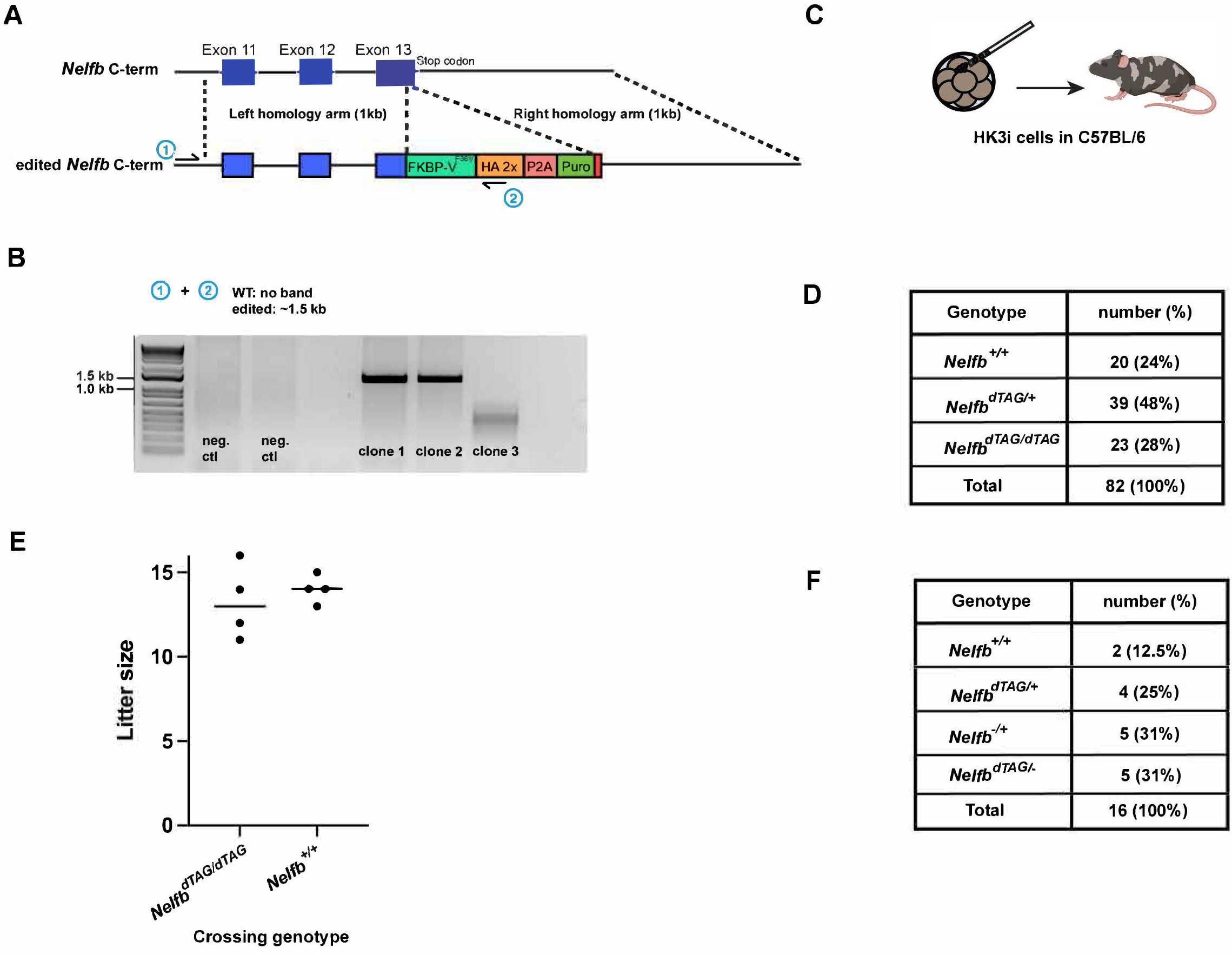
Genetic targeting and mouse generation strategies, related to Figure 1. (A) *Nelfb^dTAG^* knock-in strategy. CRISPR-Cas9 was used to generate a double-stranded break a few bases before the stop codon. The HDR template incorporates the insert immediately before the stop codon. (B) PCR validation of targeted and expanded mESC clones. Primers targets are shown in A and sequence is available in Table S2. (C) Strategy of chimeric mice generation. Further details are available in the methods section. (D)Resulting genotypes from heterozygous *NelfbdTAG/+* crossing show mendelian ratios of expected genotypes. The data suggests that the edited allele is functional. (E) Litter sizes from homozygous *Nelfb^dTAG/dTAG^* crossings are normal. (F) Resulting genotypes from heterozygous *NelfbdTAG/+ x Nelfb^+/-^* mouse suggest that the edited allele complements a null Nelfb allele.

**Figure S2.**
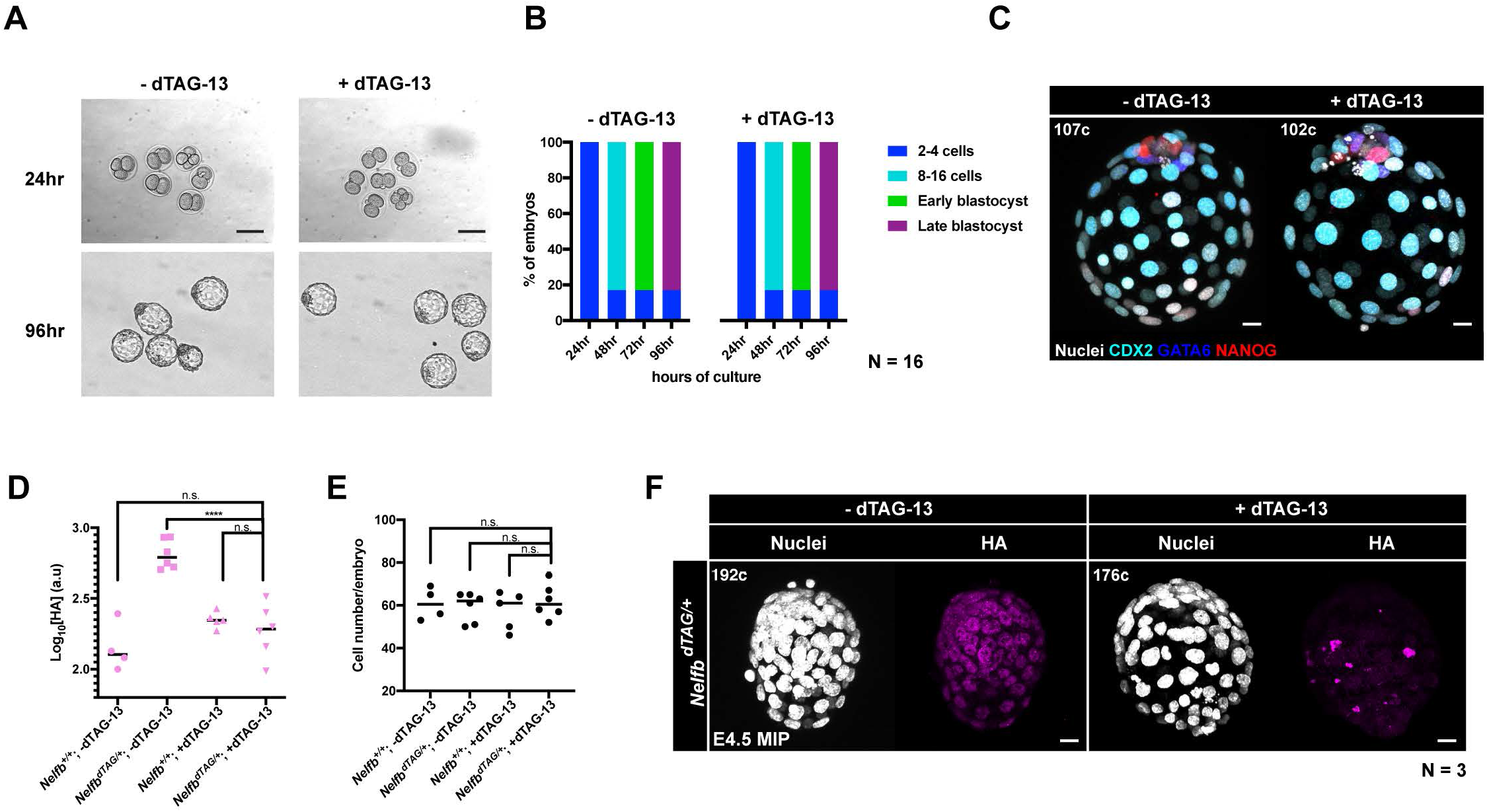
dTAG-13 is safe during pre-implantation stages, related to Figure 2. (A) Widefield images of zygotes ex utero culture +/- dTAG-13. Scale bar, 50um. (B) Percentages of cultured zygotes passing each developmental stage +/- dTAG-13. (C) Immunofluorescence of 96hr blastocysts post zygote culture +/- dTAG-13. NANOG marks the epiblast, GATA6 marks the primitive endoderm, and CDX2 marks the trophectoderm lineage. Nuclei were labeled with Hoechst. Scale bar, 50μm. (D)Quantification of mean HA signal of nuclei/embryo. Same data presented in Figure 2C but averaged per embryo. (E) Average number of nuclei/embryo post culture +/-dTAG-13 for 1hr. Cell number was determined by counting nuclei stained with Hoechst. (F) Immunofluorescence of E4.5 embryos after 1hr culture +/- dTAG-13. Nuclei are labeled with Hoechst. Total cell count per embryo is shown in the top left corner. Scale bar, 15μm. For all experiments, maximum intensity projection (MIP) is shown in images. Plots show each data point with groups means. Student t-test was used to determine significance. Statistical significance is classified based on p-value as: n.s. > 0.05, * < 0.05, ** < 0.01, *** < 0.001, **** < 0.0001.

**Figure S3.**
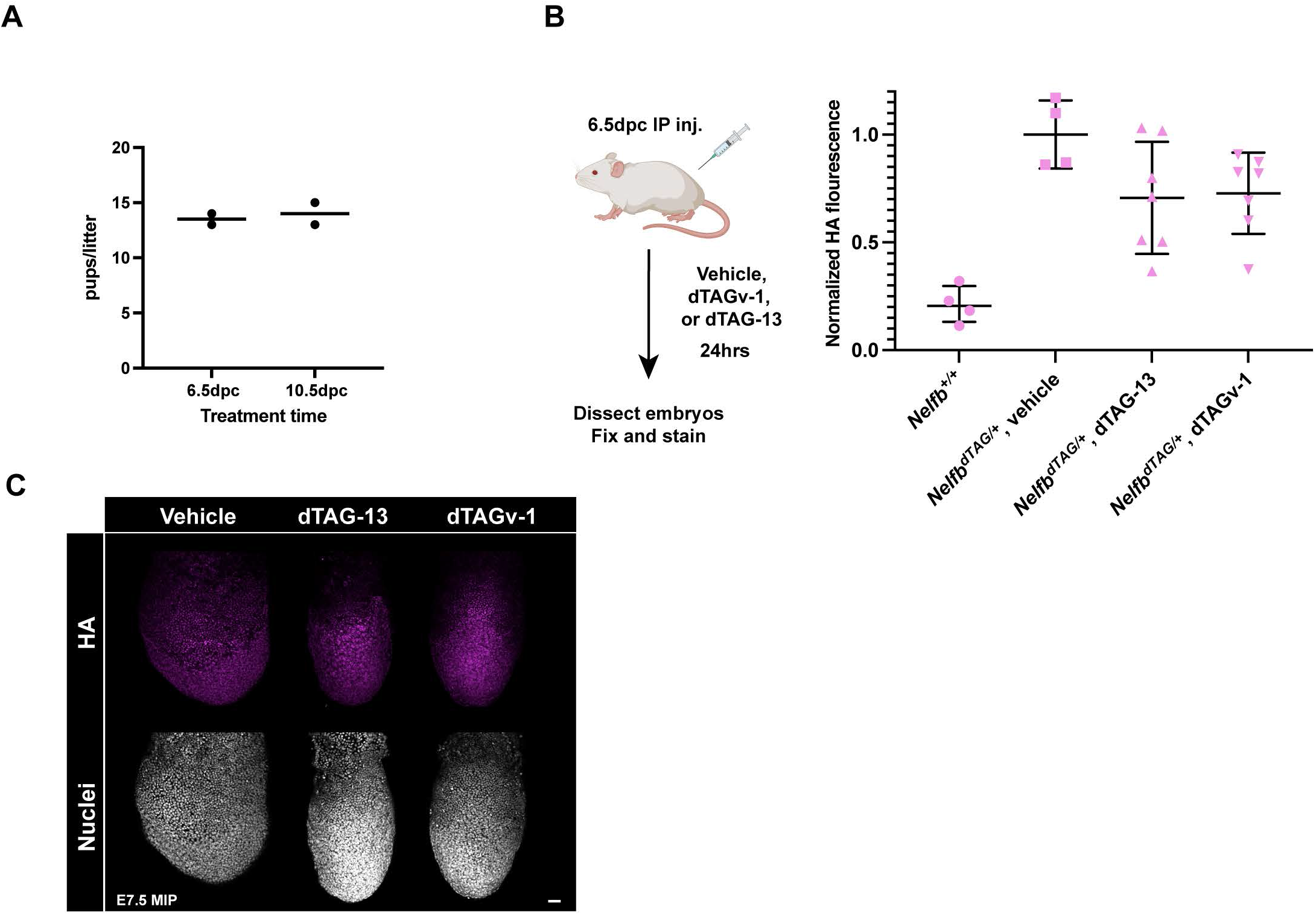
Safety and reversibility of dTAG-13 and dTAGv-1 in vivo, related to Figure 3. (A) Number of pups delivered by pregnant females after 35mg kg^-1^ dTAG-13 injection at the indicated time points. Bar represents average of two litters. (B) (Left) Schematic of the experiment. *NelfbdTAG/+* pregnant females were injected with dTAGv-1 or dTAG-13 6.5dpc. Embryos were collected for analysis 24hrs later. Quantification of mean HA signal intensity in recovered embryos 24 hr post dTAGv-1 or dTAG-13 inj. (C) Immunofluorescence images of E7.5 embryos 24hrs post dTAGv-1 or dTAG-13 injection to 6.5dpc pregnant females. Nuclei are labeled with Hoechst. Scale bar, 50μm. For all experiments, maximum intensity projection (MIP) is shown in images. Plots show each data point with groups means and interquartile range.

**Figure S4.**
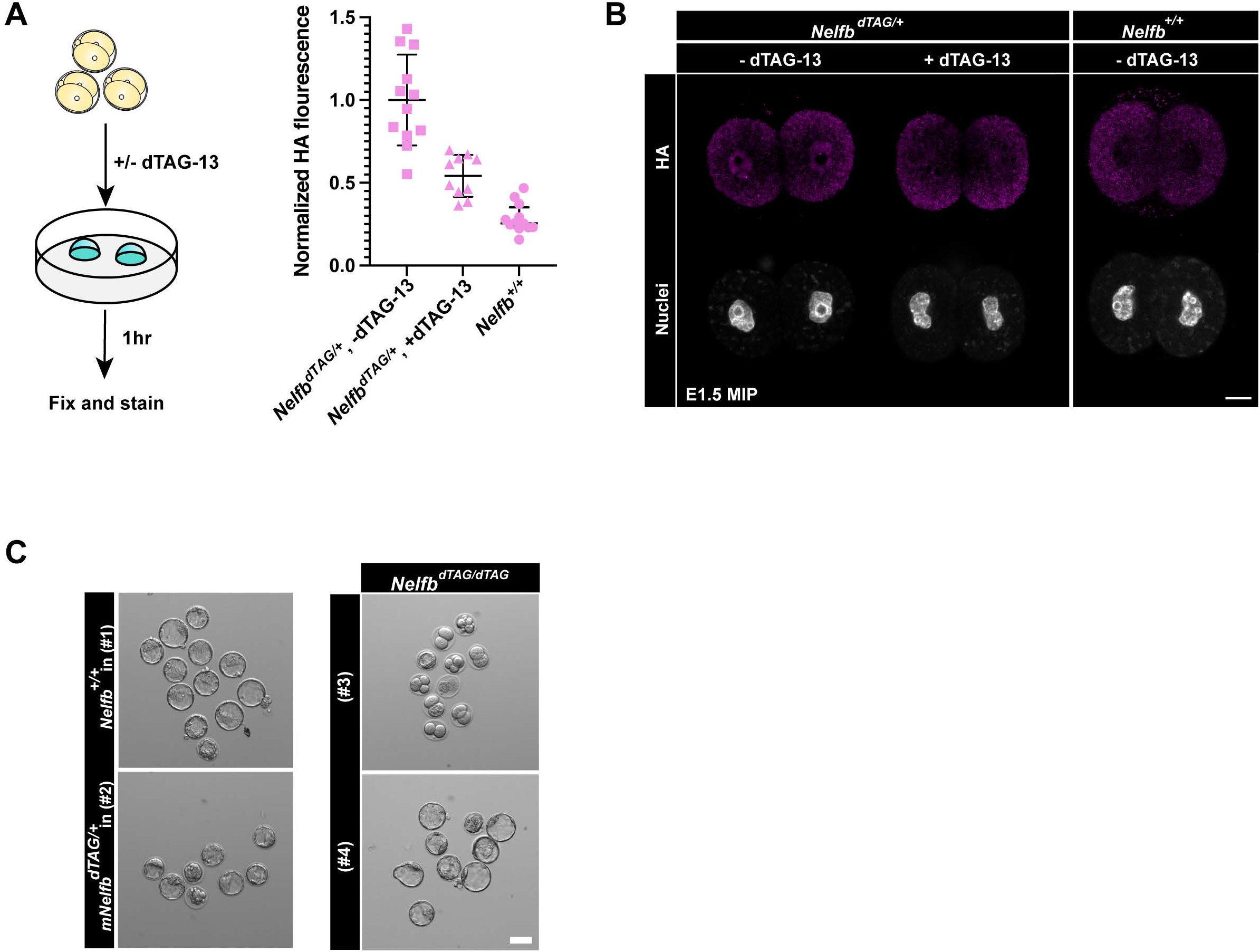
NELFB is required for pre-implantation development, related to Figure 4. (A) (Left) Schematic of the experiment. *NelfbdTAG/+* 2-cells stage embryos were culture +/- dTAG-13 for 1hr followed by immunostaining analysis. Quantification of mean HA signal in 2-cell stage embryos post 1hr culture +/- dTAG-13. Data points are single nuclei. (B) Immunofluorescence images of 2-cell stage embryos +/- dTAG-13 for 1hr. Nuclei are labeled with Hoechst. Scale bar, 10μm. (C) Additional representative images of zygote to blastocyst stage culture in Figure 4D and 4E. The treatment numbers: (#1): no dTAG-13, (#2) constant dTAG-13, (#3) 36hrs dTAG-13 from zygote to 4-cell stage followed by washing, (#4) 60hrs dTAG-13 from 4- cell stage to late blastocyst stage. Scale bar, 50μm. For all experiments, Hoechst labels nuclei. Maximum intensity projection (MIP) is shown. Plots show each data point with means.

**Figure S5.**
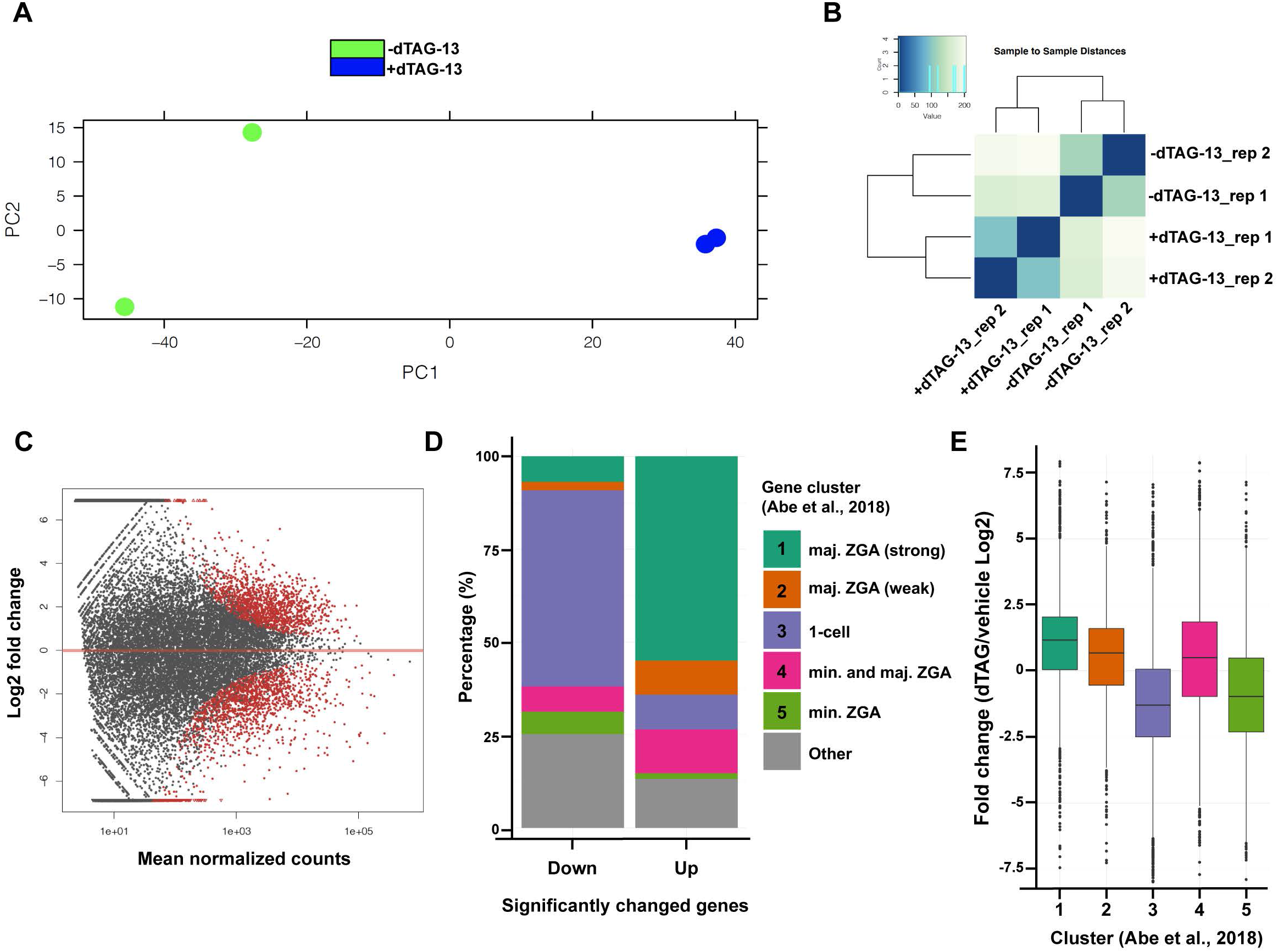
Clustering of RNA-seq samples and identity of differentially expressed genes, related to Figure 5. (A) PCA plot of RNA-seq replicates. (B) Sample-to-sample distance analysis of RNA-seq replicates. (C) MA plot of gene expression levels (+dTAG-13/-dTAG-13) from DEseq2. (D)Classes of up and downregulated genes using Abe et al., 2018 clustering. (E) Overall expression changes in clusters in Abe et al., 2018.

## REFERENCES

Abe, K.-I., Funaya, S., Tsukioka, D., Kawamura, M., Suzuki, Y., Suzuki, M.G., Schultz, R.M., and Aoki, F. (2018). Minor zygotic gene activation is essential for mouse preimplantation development. Proc. Natl. Acad. Sci. U. S. A. 115, E6780–E6788.

Adelman, K., and Lis, J.T. (2012). Promoter-proximal pausing of RNA polymerase II: emerging roles in metazoans. Nat. Rev. Genet. 13, 720–731.

Amleh, A., Nair, S.J., Sun, J., Sutherland, A., Hasty, P., and Li, R. (2009). Mouse cofactor of BRCA1 (Cobra1) is required for early embryogenesis. PLoS One 4, 2–9.

Bardot, E.S., and Hadjantonakis, A.-K. (2020). Mouse gastrulation: Coordination of tissue patterning, specification and diversification of cell fate. Mech. Dev. 163, 103617.

Behringer, R., Gertsenstein, M., Nagy, K.V., and Nagy, A. (2014). Manipulating the Mouse Embryo: A Laboratory Manual (Cold Spring Harbor Laboratory Press).

Chamberlain, P.P., D’Agostino, L.A., Ellis, J.M., Hansen, J.D., Matyskiela, M.E., McDonald, J.J., Riggs, J.R., and Hamann, L.G. (2019). Evolution of Cereblon-Mediated Protein Degradation as a Therapeutic Modality. ACS Med. Chem. Lett. 10, 1592–1602.

Clift, D., McEwan, W.A., Labzin, L.I., Konieczny, V., Mogessie, B., James, L.C., and Schuh, M. (2017). A Method for the Acute and Rapid Degradation of Endogenous Proteins. Cell 171, 1692–1706.e18.

Core, L., and Adelman, K. (2019). Promoter-proximal pausing of RNA polymerase II: a nexus of gene regulation. Genes Dev. 33, 960–982.

Hu, Z., Tan, D.E.K., Chia, G., Tan, H., Leong, H.F., Chen, B.J., Lau, M.S., Tan, K.Y.S., Bi, X., Yang, D., et al. (2020). Maternal factor NELFA drives a 2C-like state in mouse embryonic stem cells. Nat. Cell Biol. 22, 175–186.

Jaeger, M.G., and Winter, G.E. (2021). Fast-acting chemical tools to delineate causality in transcriptional control. Mol. Cell 81, 1617–1630.

Kiyonari, H., Kaneko, M., Abe, S., and Aizawa, S. (2010). Three inhibitors of FGF receptor, ERK, and GSK3 establishes germline-competent embryonic stem cells of C57BL/6N mouse strain with high efficiency and stability. Genesis 48, 317–327.

Kwak, H., and Lis, J.T. (2013). Control of Transcriptional Elongation. Annu. Rev. Genet. 47, 483–508.

Li, W., Germain, R.N., and Gerner, M.Y. (2019). High-dimensional cell-level analysis of tissues with Ce3D multiplex volume imaging. Nat. Protoc. 14, 1708–1733.

Liu, B., Xu, Q., Wang, Q., Feng, S., Lai, F., Wang, P., Zheng, F., Xiang, Y., Wu, J., Nie, J., et al. (2020). The landscape of RNA Pol II binding reveals a stepwise transition during ZGA. Nature 587, 139–144.

Lou, X., Kang, M., Xenopoulos, P., Munoz-Descalzo, S., and Hadjantonakis, A.-K. (2014). A rapid and efficient 2D/3D nuclear segmentation method for analysis of early mouse embryo and stem cell image data. Stem Cell Reports 2, 382–397.

Love, M.I., Huber, W., and Anders, S. (2014). Moderated estimation of fold change and dispersion for RNA-seq data with DESeq2. Genome Biol. 15, 550.

Nabet, B., Roberts, J.M., Buckley, D.L., Paulk, J., Dastjerdi, S., Yang, A., Leggett, A.L., Erb, M.A., Lawlor, M.A., Souza, A., et al. (2018). The dTAG system for immediate and target-specific protein degradation. Nat. Chem. Biol. 14, 431–441.

Nabet, B., Ferguson, F.M., Seong, B.K.A., Kuljanin, M., Leggett, A.L., Mohardt, M.L., Robichaud, A., Conway, A.S., Buckley, D.L., Mancias, J.D., et al. (2020). Rapid and direct control of target protein levels with VHL-recruiting dTAG molecules. Nat. Commun. 11, 4687.

Nishimura, K., Fukagawa, T., Takisawa, H., Kakimoto, T., and Kanemaki, M. (2009). An auxin-based degron system for the rapid depletion of proteins in nonplant cells. Nat. Methods 6, 917–922.

Nishimura, K., Yamada, R., Hagihara, S., Iwasaki, R., Uchida, N., Kamura, T., Takahashi, K., Torii, K.U., and Fukagawa, T. (2020). A super-sensitive auxin-inducible degron system with an engineered auxin-TIR1 pair. Nucleic Acids Res. 48, e108.

Nowotschin, S., Setty, M., Kuo, Y.-Y., Liu, V., Garg, V., Sharma, R., Simon, C.S., Saiz, N., Gardner, R., Boutet, S.C., et al. (2019). The emergent landscape of the mouse gut endoderm at single-cell resolution. Nature 569, 361–367.

Pickar-Oliver, A., and Gersbach, C.A. (2019). The next generation of CRISPR-Cas technologies and applications. Nat. Rev. Mol. Cell Biol. 20, 490–507.

Pijuan-Sala, B., Griffiths, J.A., Guibentif, C., Hiscock, T.W., Jawaid, W., Calero-Nieto, F.J., Mulas, C., Ibarra-Soria, X., Tyser, R.C. V, Ho, D.L.L., et al. (2019). A single-cell molecular map of mouse gastrulation and early organogenesis. Nature 566, 490–495.

Piliszek, A., Kwon, G.S., and Hadjantonakis, A.-K. (2011). Ex utero culture and live imaging of mouse embryos. Methods Mol. Biol. 770, 243–257.

Pimeisl, I.-M., Tanriver, Y., Daza, R.A., Vauti, F., Hevner, R.F., Arnold, H.-H., and Arnold, S.J. (2013). Generation and characterization of a tamoxifen-inducible Eomes(CreER) mouse line. Genesis 51, 725–733.

Poueymirou, W.T., Auerbach, W., Frendewey, D., Hickey, J.F., Escaravage, J.M., Esau, L., Doré, A.T., Stevens, S., Adams, N.C., Dominguez, M.G., et al. (2007). F0 generation mice fully derived from gene-targeted embryonic stem cells allowing immediate phenotypic analyses. Nat. Biotechnol. 25, 91–99.

Ran, F.A., Hsu, P.D., Wright, J., Agarwala, V., Scott, D.A., and Zhang, F. (2013). Genome engineering using the CRISPR-Cas9 system. Nat. Protoc. 8, 2281–2308.

Saiz, N., Plusa, B., and Hadjantonakis, A.-K. (2015). Single cells get together: High-resolution approaches to study the dynamics of early mouse development. Semin. Cell Dev. Biol. 47–48, 92– 100.

Saiz, N., Williams, K.M., Seshan, V.E., and Hadjantonakis, A.-K. (2016). Asynchronous fate decisions by single cells collectively ensure consistent lineage composition in the mouse blastocyst. Nat. Commun. 7, 13463.

Savery, D., Maniou, E., Culshaw, L.H., Greene, N.D.E., Copp, A.J., and Galea, G.L. (2020). Refinement of inducible gene deletion in embryos of pregnant mice. Birth Defects Res. 112, 196–204.

Schindelin, J., Arganda-Carreras, I., Frise, E., Kaynig, V., Longair, M., Pietzsch, T., Preibisch, S., Rueden, C., Saalfeld, S., Schmid, B., et al. (2012). Fiji: an open-source platform for biological-image analysis. Nat. Methods 9, 676–682.

Schrode, N., Xenopoulos, P., Piliszek, A., Frankenberg, S., Plusa, B., and Hadjantonakis, A.-K. (2013). Anatomy of a blastocyst: Cell behaviors driving cell fate choice and morphogenesis in the early mouse embryo. Genesis 51, 219–233.

Smits, A.H., Ziebell, F., Joberty, G., Zinn, N., Mueller, W.F., Clauder-Münster, S., Eberhard, D., Fälth Savitski, M., Grandi, P., Jakob, P., et al. (2019). Biological plasticity rescues target activity in CRISPR knock outs. Nat. Methods 16, 1087–1093.

Sun, Q.-Y., Liu, K., and Kikuchi, K. (2008). Oocyte-specific knockout: a novel in vivo approach for studying gene functions during folliculogenesis, oocyte maturation, fertilization, and embryogenesis. Biol. Reprod. 79, 1014–1020.

Tadros, W., and Lipshitz, H.D. (2009). The maternal-to-zygotic transition: a play in two acts. Development 136, 3033–3042.

Ved, N., Curran, A., Ashcroft, F.M., and Sparrow, D.B. (2019). Tamoxifen administration in pregnant mice can be deleterious to both mother and embryo. Lab. Anim. 53, 630–633.

Verma, R., Mohl, D., and Deshaies, R.J. (2020). Harnessing the Power of Proteolysis for Targeted Protein Inactivation. Mol. Cell 77, 446–460.

Vos, S.M., Farnung, L., Urlaub, H., and Cramer, P. (2018a). Structure of paused transcription complex Pol II–DSIF–NELF. Nature.

Vos, S.M., Farnung, L., Urlaub, H., and Cramer, P. (2018b). Structure of paused transcription complex Pol II–DSIF–NELF. Nature 560, 601–606.

Vos, S.M., Farnung, L., Boehning, M., Wigge, C., Linden, A., Urlaub, H., and Cramer, P. (2018c). Structure of activated transcription complex Pol II-DSIF-PAF-SPT6. Nature 560, 607–612.

Wang, X., Hang, S., Prazak, L., and Gergen, J.P. (2010). NELF Potentiates Gene Transcription in the Drosophila Embryo. PLoS One 5, e11498.

Williams, L.H., Fromm, G., Gokey, N.G., Henriques, T., Muse, G.W., Burkholder, A., Fargo, D.C., Hu, G., and Adelman, K. (2015). Pausing of RNA Polymerase II Regulates Mammalian Developmental Potential through Control of Signaling Networks. Mol. Cell 58, 311–322.

Wu, T., Yoon, H., Xiong, Y., Dixon-Clarke, S.E., Nowak, R.P., and Fischer, E.S. (2020). Targeted protein degradation as a powerful research tool in basic biology and drug target discovery. Nat. Struct. Mol. Biol. 27, 605–614.

Yamaguchi, Y., Takagi, T., Wada, T., Yano, K., Furuya, A., Sugimoto, S., Hasegawa, J., and Handa, H. (1999). NELF, a multisubunit complex containing RD, cooperates with DSIF to repress RNA polymerase II elongation. Cell 97, 41–51.

Yesbolatova, A., Saito, Y., Kitamoto, N., Makino-Itou, H., Ajima, R., Nakano, R., Nakaoka, H., Fukui, K., Gamo, K., Tominari, Y., et al. (2020). The auxin-inducible degron 2 technology provides sharp degradation control in yeast, mammalian cells, and mice. Nat. Commun. 11, 5701.

